# Functional metabolic phenotyping of human pancreatic ductal adenocarcinoma

**DOI:** 10.1101/2021.07.23.452145

**Authors:** Irina Heid, Sinan Karakaya, Corinna Münch, Smiths S. Lueong, Alina M. Winkelkotte, Sven T. Liffers, Laura Godfrey, Phyllis FY Cheung, Konstatinos Savvatakis, Geoffrey J. Topping, Florian Englert, Lukas Kritzner, Martin Grashei, Thomas Kunzke, Na Sun, Axel Walch, Andrea Tannapfel, Richard Viebahn, Heiner Wolters, Waldemar Uhl, Deepak Vangala, Esther M.M. Smeets, Erik H.J.G. Aarntzen, Daniel Rauh, Jörg D. Hoheisel, Doris Hellerschmied, Stephan A. Hahn, Franz Schilling, Rickmer Braren, Marija Trajkovic-Arsic, Jens T. Siveke

## Abstract

Pancreatic Ductal Adenocarcinoma (PDAC) lacks targeted treatment options. Although subtypes with transcriptome-based distinct lineage and differentiation features have been identified, deduced clinically actionable targets remain elusive. We here investigate functional metabolic features of the classical and QM (quasi-mesenchymal)/basal-like PDAC subtypes potentially exploitable for non-invasive subtype differentiation and therapeutic intervention.

A collection of human PDAC cell lines, primary patient derived cells (PDC), patient derived xenografts (PDX) and patient PDAC samples were transcriptionally stratified into the classical and QM subtype. Functional metabolic analyses including targeted and non-targeted metabolite profiling (matrix-assisted laser desorption/ionization mass spectrometry imaging (MALDI-MSI)), seahorse metabolic flux assays and metabolic drug targeting were performed. Hyperpolarized ^13^C-magnetic resonance spectroscopy (HP-MRS) of PDAC xenografts was used for *in vivo* detection of intra-tumoral [1-^13^C]pyruvate and [1-^13^C]lactate metabolism.

We identified glycolysis and lipid metabolism/fatty acid oxidation as transcriptionally preserved metabolic pathways in QM and classical PDAC subtype respectively. However, these metabolic cues were not unambiguously functionally linked to one subtype. Striking functional metabolic heterogeneity was observed especially in primary patient derived cells with only individual samples representing high dependence on glycolysis or mitochondrial oxidation. Of note, QM cells actively use the glycolytic product lactate as oxidative mitochondrial fuel. Using HP-MRS, we were able to non-invasively differentiate glycolytic tumor xenografts with high intratumoral [1-^13^C]pyruvate to [1-^13^C]lactate conversion *in vivo*.

Although PDAC transcriptomes indicate molecular subtype-associated distinct metabolic pathways, we found substantial functional metabolic heterogeneity independent of the molecular subtype. Non-invasive identification of highly glycolytic tumors by [1-^13^C]pyruvate/lactate HP-MRS support individualized metabolic targeting approaches.

## Introduction

Despite enormous research efforts in the last 50 years, pancreatic ductal adenocarcinoma (PDAC) remains a fatal disease with marginal clinical advancement [Aung et al., 2018]. Although the oncogenic drivers as well as transcriptional and molecular profiles of PDAC have been studied in great detail [Chan-Seng-Yue et al., 2020; Moffitt et al., 2015; Waddell et al., 2015], effective targeting strategies remain scarce. Sequencing efforts in large patient cohorts have identified distinct molecular PDAC subtypes with two dominant lineages: classical/pancreatic progenitor and quasi-mesenchymal (QM) /squamous/basal-like [Aung et al., 2018; Bailey et al., 2016; Cancer Genome Atlas Research Network. Electronic address and Cancer Genome Atlas Research, 2017; Collisson et al., 2011]. QM PDACs are associated with shorter median survival and resistance to first-line chemotherapy with FOLFIRINOX [Aung et al., 2018]. Yet, which cancer cell features contribute to the aggressive and therapy-resistant phenotype phenotype remains unknown.

Metabolic plasticity, i.e. an individual cells ability to use different metabolic pathways in dependence of alternating growth conditions including oxygen and nutrient availability has been implicated as a major cause of therapy resistance in cancers [DeBerardinis and Chandel, 2016]. This metabolic plasticity allows PDAC cells not only to adapt but to thrive on particularly scarce conditions of hypoxia and nutrient limitations [Biancur and Kimmelman, 2018] typically observed in PDAC. Recent transcriptional metabolic profiling of 33 cancer entities identified seven metabolic super-pathways that are selectively altered in specific cancer subpopulations and dramatically influence sensitivity to therapy. Cancers with upregulated gene signatures for carbohydrate, nucleotide and vitamin/cofactor metabolism show worse prognosis than those with enhanced lipid metabolism [Peng et al., 2018]. In PDAC, metabolic transcripts involved in glycolysis and cholesterol biosynthesis are associated with the classical and QM subtypes, respectively [Karasinska et al., 2020]. However, functional evidence that these pathways are indeed significantly operable in defined PDAC subtypes and thus therapeutically targetable are still largely missing.

In this work, we analyzed metabolic transcripts present in the classical and QM PDAC subtypes in a large collection of samples reaching from long-term cultured PDAC cell lines to patient-derived primary model systems. We address to which extent are those transcriptomic signatures functionally mirrored and whether differences in the metabolic phenotype between subtypes allows non-invasive subtype identification. We observed strong heterogeneity in the metabolic behavior especially in patient-derived models and were able to *in vivo* non-invasively detect highly glycolytic PDACs based on high conversion of [1-^13^C]pyruvate to [1-^13^C]lactate and vice-versa by HP-MRS. Our work opens a perspective for a non-invasive monitoring of personalized metabolic targeting approaches.

## Results

### Glycolysis and lipid metabolic transcripts are preserved in the classical and QM PDAC subtypes

To analyze which metabolic features are associated with molecular PDAC subtypes, we first performed transcriptome-based molecular subtyping in multiple preclinical and clinical samples. RNA was isolated from conventional PDAC cell lines (n=8), patient derived xenografts (PDX, n=34) and primary patient derived cells (PDC, n=11) and RNA-seq or Microarray analysis was performed. Transcriptomes from bulk tissue of 204 PDAC samples from previously published resource was utilized (E-MTAB-1791).

For tumor subtype determination, we used publicly available transcriptionally subtyped PDAC cohorts (PDAC cell lines (GSE21654 [Maupin et al., 2010]) PDAC xenograft (E-MTAB-4029 [Noll et al., 2016]) and bulk PDAC tissue (GSE16515 and GSE15471 [Pei et al., 2009] [Badea et al., 2008]) for benchmarking. After that, patient PDAC samples, PDX cohort and PDC samples were stratified to QM and classical group and gene set enrichment analysis (GSEA) was performed. In the PDAC patient sample cohort (204 samples), 88 were classified as classical and 116 as QM subtype. Samples clustering to the QM subtype presented significant enrichment of selected QM and squamous subtype assigner gene sets previously described [Bailey et al., 2016; Collisson et al., 2011] (figure 1b; supplementary table 1) supporting correct subtype assignment.

**Figure 1:**
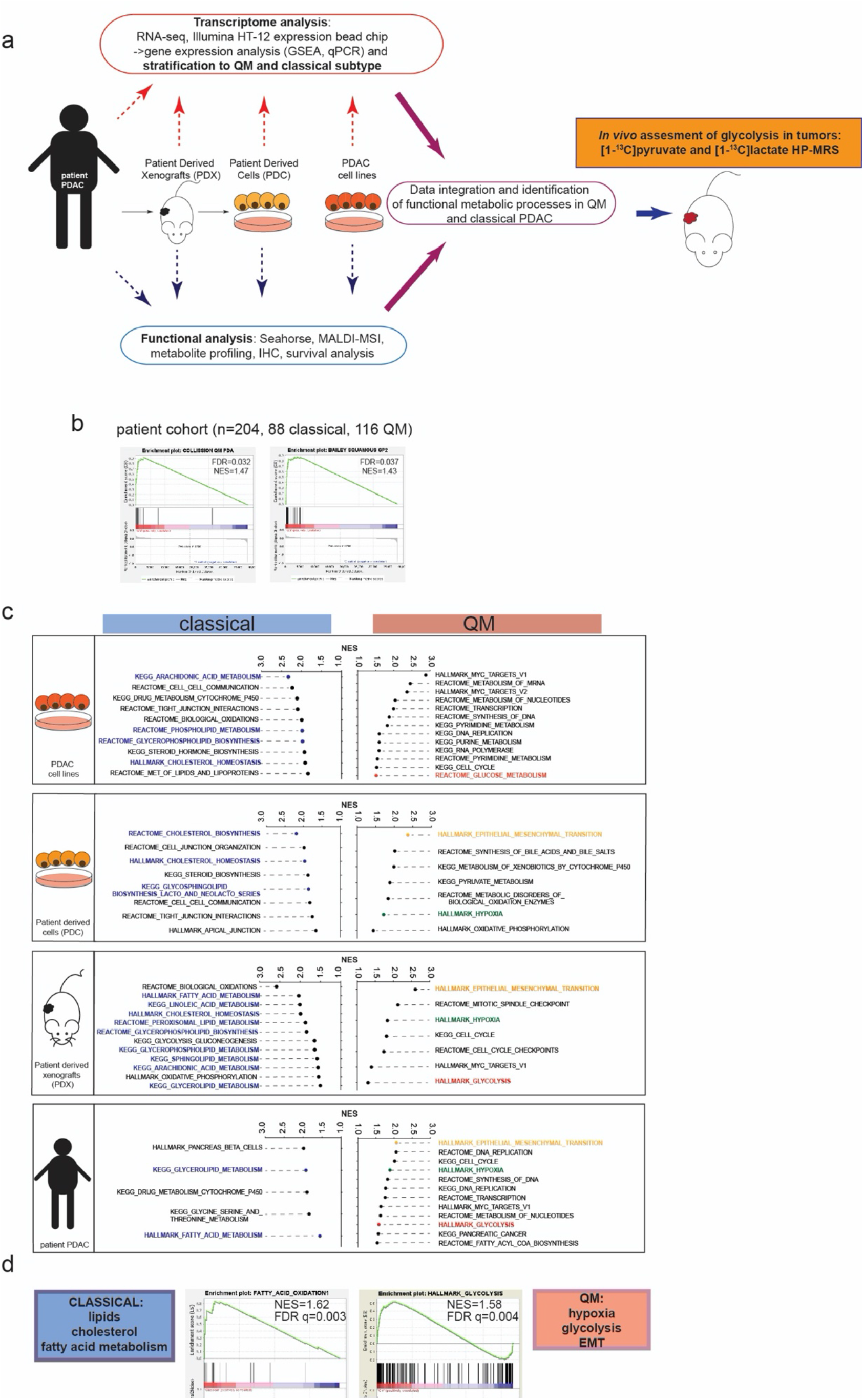
Gene Set Enrichment Analysis (GSEA) identifies glycolysis, hypoxia and lipid/fatty acid metabolism are enriched in QM and classical PDAC samples respectively. a) Graphical sketch of the used models and experimental flow in the study. b) Enrichment plots for the selected “Collisson QM” and “Bailey squamous GP2” assigner gene sets in our patient cohort. Both gene sets are enriched in here defined QM PDAC samples. FDR and NES presented in the figure. c) GSEA analysis for QM vs classical groups was performed for cell lines (n=8; 4QM, 4 classical), Patient Derived Cells (PDC; n=11, 5 QM and 6 classical) Patient Derived Xenografts (PDX; n=34, 12 QM and 22 classical), and patient PDAC samples (n=204; 116 QM, 88 classical). Presented are Normalized Enrichment Scores (NES) values for selection of metabolic gene sets identified as significantly enriched (False Discovery Rate, FDR q value <0.06) in QM or classical subtypes. Gene set databases HALLMARK, REACTOME and KEGG were used for analysis. Epithelial-to-mesenchymal transition (EMT, blue), glycolysis/glucose metabolism (orange), hypoxia (green) and MYC targets gene sets are commonly enriched in most of the QM datasets. In classical subtype, gene sets typical for cellular organization (tight junctions, cell-cell communication) together with lipid/cholesterol/fatty acid metabolism (dark blue) are enriched. d) Enrichment plots for Fatty Acid Oxidation (FAO) generated gene set and HALLMARK glycolysis gene set specifically enriched in classical and QM PDAC patient samples respectively.

In the PDX cohort, 22 classical and 12 QM tumors were identified and 6 classical and 5 QM among 11 PDCs. The 8 PDAC cell lines used in this study were previously classified as QM (KP4, PSN1, MIAPaca2, PaTu8988T) and classical (PaTu8988S, HUPT4, HPAFII, HPAC) [Daemen et al., 2015]. We analyzed gene expression of Vimentin (*VIM*) and E-cadherin (*CDH1*) as markers of mesenchymal and epithelial status respectively. QM PDAC cell lines presented high *VIM* and low *CDH1* gene expression as typical for mesenchymal feature enrichment. Classical PDAC cell lines presented higher CDH1 and lower VIM expression (supplementary figure 1a). *VIM* and *CDH1* expression correlated well with the subtype of the PDCs as well (supplementary figure 1b).

After classification, QM and classical groups were compared by GSEA for HALLMARK, REACTOME and KEGG collections in all datasets. A full list of all enriched gene sets with respective Normalized Enrichment Score (NES) and False Discovery Rate (FDR) values is given in supplementary table 2. As expected for the mesenchymal phenotype, enrichment of epithelial-to-mesenchymal transition (EMT) gene set was observed in the QM group in PDX, PDC and patient PDAC samples (figure 1c). Analysis of metabolic transcripts revealed a remarkable stability of subtype-typical metabolic pathways throughout different models (figure 1c). Transcripts involved in lipid metabolism (glycerophospholipid, sphingolipid, glycerolipid, glycolipid) as well as cholesterol metabolism were generally enriched in classical samples. In PDX and patient PDAC samples of the classical subtype, the fatty acid (FA) metabolism gene set was also strongly enriched, suggesting not only structural but also active metabolic role of lipids in the classical subtype. Therefore, we next analyzed a generated fatty acid oxidation gene set containing 14 genes involved exclusively in the mitochondrial beta oxidation (FAO1) (supplementary table 3). This gene set was also significantly enriched in the classical patient PDAC samples (Figure 1d).

In QM samples, transcripts involved in glycolysis and hypoxia were preserved (figure 1c). The hypoxia gene set was enriched in QM bulk PDAC tissue, PDX and PDC data sets, even though PDC cells were cultured under common laboratory normoxic conditions. Glycolysis/glucose metabolism as well as MYC-targets gene sets were also enriched in the QM patient PDAC samples, PDX and PDAC cell lines datasets. Interestingly, the glycolysis gene set was not enriched in the QM PDCs, possibly due to low sample numbers but also suggesting no unambiguous assignment of glycolytic genes to the QM subtype at least in PDCs. In summary, we observed strong transcriptional association of classical and QM subtypes with lipid/FA metabolism and glycolysis respectively.

### Classical and QM PDACs differ in lipid metabolism

To address whether the identified metabolic transcripts are effectively translated into active lipid and glucose metabolism in the classical and QM subtype respectively, we first analyzed distribution of structural lipids, energy storing lipids and free fatty acid in PDAC cell lines and primary PDCs.

Targeted metabolite profiling revealed enrichment of different structural lipids (sphingomyelins, lysophosphatidylcholines, phosphatidylcholines) in the classical PDAC cell lines (figure 2a, supplementary table 4) similar to what was previously described [Daemen et al., 2015]. In PDCs, a more heterogeneous distribution pattern was observed with generally higher accumulation of some structural lipids in the classical PDCs, however not as pronounced as in PDAC cell lines (figure 2a). These observations may hint for differences in the management of structural lipids in classical and QM subtypes.

**Figure 2:**
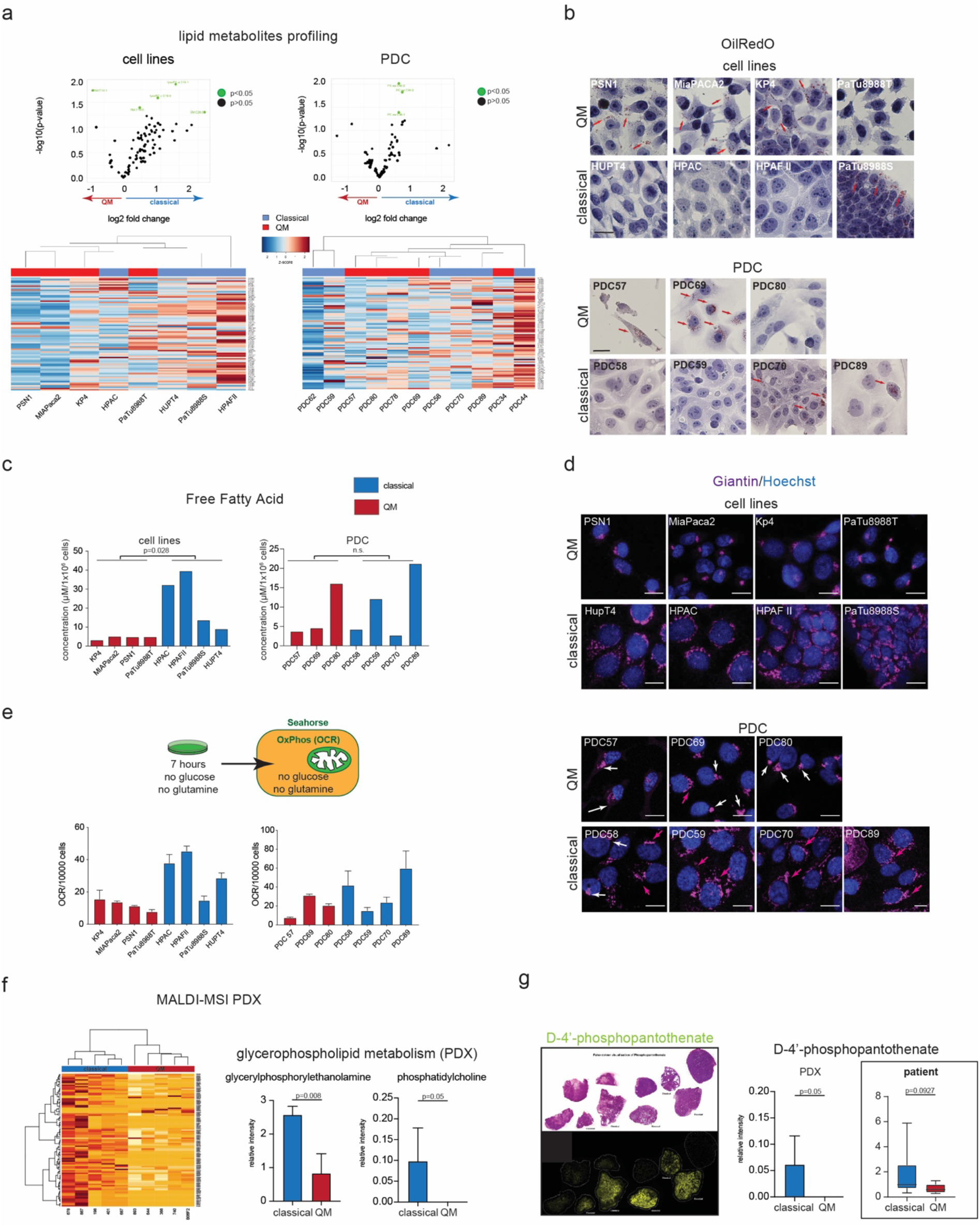
Differences in lipid metabolism in QM and classical PDACs. a) Volcano plots and hierarchical clustering for lipid metabolites, PC-phospatidylcholines, LPC-lysophospatidilcholines and SM-sphingomyelins in the QM and classical PDAC cell lines and PDCs as measured by Biocrates Absolute p180 kit. Upper panel: Volcano plots showing general enrichment of lipid metabolites in classical cell lines. This effect was present but less pronounced in PDCs. One dot presents one metabolite. Green dots present significantly changed metabolites between classical and QM subtype with p <0.05 (Student’s T-test, unpaired, two-sided). Full list of metabolites, measured concentrations and abbreviations is given in supplementary table 4. Lower panels: hierarchical clustering of all analyzed structural lipid metabolites according to their measured concentrations in PDAC cell lines (84 metabolites) and PDCs (85 metabolites). Z score: red color indicates high, blue low intensity; b) OilRedO staining for lipid droplets in PDAC cells. Accumulation of lipid droplets observed in QM cell lines PSN1, MIAPaCa2, Kp4, in PDC69 and occasionally in PDC57. Among classical cell lines PaTu8988S was very rich in lipid droplets while in HPAFII, HPAC and HUPT4 cells lipid droplets were not detected. Classical PDC70 readily presented OilRed positive cells as well, while in PDC89 only very occasional single cells were positive. In classical PDC58, PDC59 lipid droplets were not detected. Red arrows indicate lipid droplets. Scale bar=10μm. c) Higher free fatty acid (FFA) in classical than in QM cell lines. PDC89 (classical) presents the highest level of FFA among PDCs. P-values calculated by the Mann-Whitney test. d) Immunofluorescence for Giantin, a Golgi membrane protein. Compact Golgi observed in QM cell lines PSN1, MIAPaca2, KP4, PaTu8988T and primary QM PDC57 and PDC80 cells. Disperse Golgi in classical lines HPAFII, HPAC, HUPT4, PaTu8988S and classical PDC59, PDC70, PDC89. Mixed Golgi structures with predominantly compact morphology observed in PDC69 (QM) and PDC58 (classical). Compact Golgi-white arrows. Disperse Golgi-red arrows. Scale bar 10μm. e) OCR levels measured for cell lines and PDCs after 7 hours of cultivation in media without glucose or glutamine. HPAC, HPAFII, HUPT4 and PDC89 present high relative OCR levels suggesting oxidation of endogenous fatty acid. Presented are OCR values (mean±SD) calculated from 2-3 wells/cell line/per 10.000 seeded cells in one experiment. f) MALDI-MSI and m/z species clustering for classical (n=5) and QM (n=5) PDX samples. Left: hierarchical clustering of differentially expressed m/z species in PDX samples. Significantly changed m/z species (Mann-Whitney test) were included in the clustering. Red color-high intensity; Light yellow-low intensity; Metabolites of glycerophospholipid metabolism, glycerylphosphorylethanolamine (m/z=214.049) and phosphatidylcholine (m/z=794.509) are significantly higher in the classical PDX samples. P-values calculated by Mann-Whitney test. g) H&E and false color visualization of D-4’-phosphopantothenate (m/z=280.0595) in cryo-sections of PDX samples. D-4’-phosphopantothenate is detected exclusively in classical PDX. D-4’-phosphopantothenate is detected exclusively in classical PDX. Framed graph: D-4’-phosphopantothenate is also prominently enriched in the human classical FFPE samples as well (classical n=9, QM n= 8). P-values calculated by Mann-Whitney test.

Next to structural lipids, storage lipids and FA are key branches of lipid metabolism. We thus investigated their distribution in 8 PDAC cell lines and 7 selected PDCs. OilRedO staining for storage lipids (neutral lipids, tryacylglycerols) revealed accumulation of lipid droplets in the QM lines PSN1, MIAPaca2 and Kp4 and in one classical line (PaTu8988S) (figure 2b). In contrast, lipid droplets were not detected in the classical cell lines HUPT4, HPAFII and HPAC nor in PaTu8988T cells. In primary PDCs, only PDC69 (QM) presented very high numbers of intracellular lipid droplets present in the majority of the cells, whereas scarce positive cells were found in PDC57 (QM). OliRedO positive cells were readily observed in classical PDC70 (30-40% of cells). In classical PDC58, PDC59 and PDC89 cells, OilRedO positive cells were in general not detected with only very few positive cells found in PDC89 (figure 2b). In terms of FA distribution, all 4 classical PDAC lines presented higher levels of free fatty acid (FFA) compared to the 4 QM lines (figure 2c). Among PDCs, the highest FFA levels were measured in classical PDC89 (figure 2c). In other investigated PDCs, FFA content did not correlate with the molecular subtype. PDC80, though QM, presented relatively high FFA levels and comparable to a classical PDC59. Taken together, QM PDAC cell lines preferably stored their FA in form of lipid droplets, while in classical cells FA was freely available for cellular processes such as FA-mitochondrial beta oxidation or incorporation into structural lipids. In PDCs, a similar trend but higher diversity in distribution of FA and storage lipids was observed.

Both structural and energy storing lipids are synthesized and matured in the endoplasmatic reticulum and the Golgi complex [Fagone and Jackowski, 2009; Pol et al., 2014]. Considering the observed differences in lipid management in QM and classical subsets, we analyzed Golgi complex morphology by anti-giantin immunofluorescence staining and observed a remarkable subtype-dependent Golgi morphology. In QM cell lines and PDCs, we observed a highly compact and well-organized Golgi complex with perinuclear localization. In contrast, classical cell lines and PDCs showed a dispersed Golgi complex (figure 2d). In some PDCs, we also observed heterogeneity within one cell population. In PDC69, though classified as QM and with a predominantly compact Golgi, some cells presented disperse Golgi structures as well. The same was true for classical PDC58 cells that presented compact and dispersed Golgi as well. In summary, Golgi complex showed a subtype-associated morphological organization potentially reflecting different needs of classical and QM cells for structural and energy lipids observed above.

Distribution of FA and energy storing lipids suggested that QM PDAC cells do not use but rather deposit the FA in lipid droplets, while classical cells have FA freely available for eventual use in mitochondrial beta-oxidation as well. We thus investigated lipid metabolism in QM and classical cells by using the seahorse metabolic flux assays (figure 2e). These real-time assays are performed in living cells and evaluate Extracellular Acidification Rate (ECAR) and Oxygen Consumption Rate (OCR) as readouts of two major energy supplying processes, glycolysis and oxidative phosphorylation (OxPhos) respectively. We designed a short-term energy evaluating seahorse experiment by cultivating the PDAC cell lines and PDCs for 7 hours in media without glucose or glutamine where only intracellular intrinsically available resources, such as FA, are present. Basal cellular OCR was then measured. In such conditions, higher OCRs were observed in HPAF II, HPAC, HUPT4 and PDC89 classical cells (figure 2e) among cell lines and PDCs respectively, potentially attributable to oxidation of intrinsically available FA. Taken together, metabolic flux assays suggest that some classical cells actively oxidize FA to maintain their basal metabolism.

To validate these findings in a more complex and translational *ex vivo* setting, we analyzed the metabolite distribution in fresh-frozen PDX tumor samples as well. For this purpose, non-targeted metabolic profiling of cryo-preserved PDX tissues using MALDI-MSI (matrix-assisted laser desorption/ionization- mass spectrometry imaging) was used. As in cells, accumulation of structural lipids in the classical PDX was detected. In 10 PDX (5 QM vs 5 classical), metabolite clustering into classical and QM groups was observed despite the limited number of samples (figure 2f). Considering significantly altered metabolites between classical and QM revealed by a U-test, we performed metabolic pathway analysis (supplementary figure 2a). Glycerophospholipid metabolism was among the top 5 most changed pathways with glycerylphosphorylethanolamine (m/z=214.049) and phosphatidylcholine (m/z=794.509) being significantly higher in the classical samples (figure 2f). Interestingly, D-4’-phosphopantothenate (m/z=280.0595), a coenzyme A (CoA) precursor, was expressed exclusively in classical PDX tumors (figure 2g). Additionally, we also performed MALDI-MSI in a cohort of human FFPE PDAC samples (tissue microarray, n=17). Samples were stratified to QM and classical based on histological expression of KRT81 and HNF1A expression as previously reported [Muckenhuber et al., 2018]. As in PDX tumors, higher levels of D-4’-phosphopantothenate were detected in the classical human PDAC FFPE samples (figure 2g). CoA is central for many enzymatic reactions in lipid synthesis and FA oxidation [Rohrig and Schulze, 2016], probably underlying the enrichment of D-4’-phosphopantothenate in the classical samples.

Taken together, prominent accumulation of structural lipids was detected in classical patient derived xenografts indicating preservation of lipid metabolic routes in a relevant patient-derived PDAC model system.

### Glycolysis is activated in selected PDAC cells

Gene set enrichment analysis pinpointed glycolysis as the most prominent metabolic pathway present in QM samples. To confirm whether glycolysis is indeed active in QM PDAC cells, we performed the seahorse metabolic flux assay and evaluated glycolysis (ECAR) and OxPhos (OCR) in cell lines and PDCs cultivated in media containing physiological concentrations of glucose (5mM) and glutamine (2mM). Under these conditions, PSN1 and PDC69, both QM, presented the highest ECAR/OCR ratios among cell lines and PDCs respectively (figure 3a), indicating higher glycolytic activity in these cells (figure 3a).

**Figure 3:**
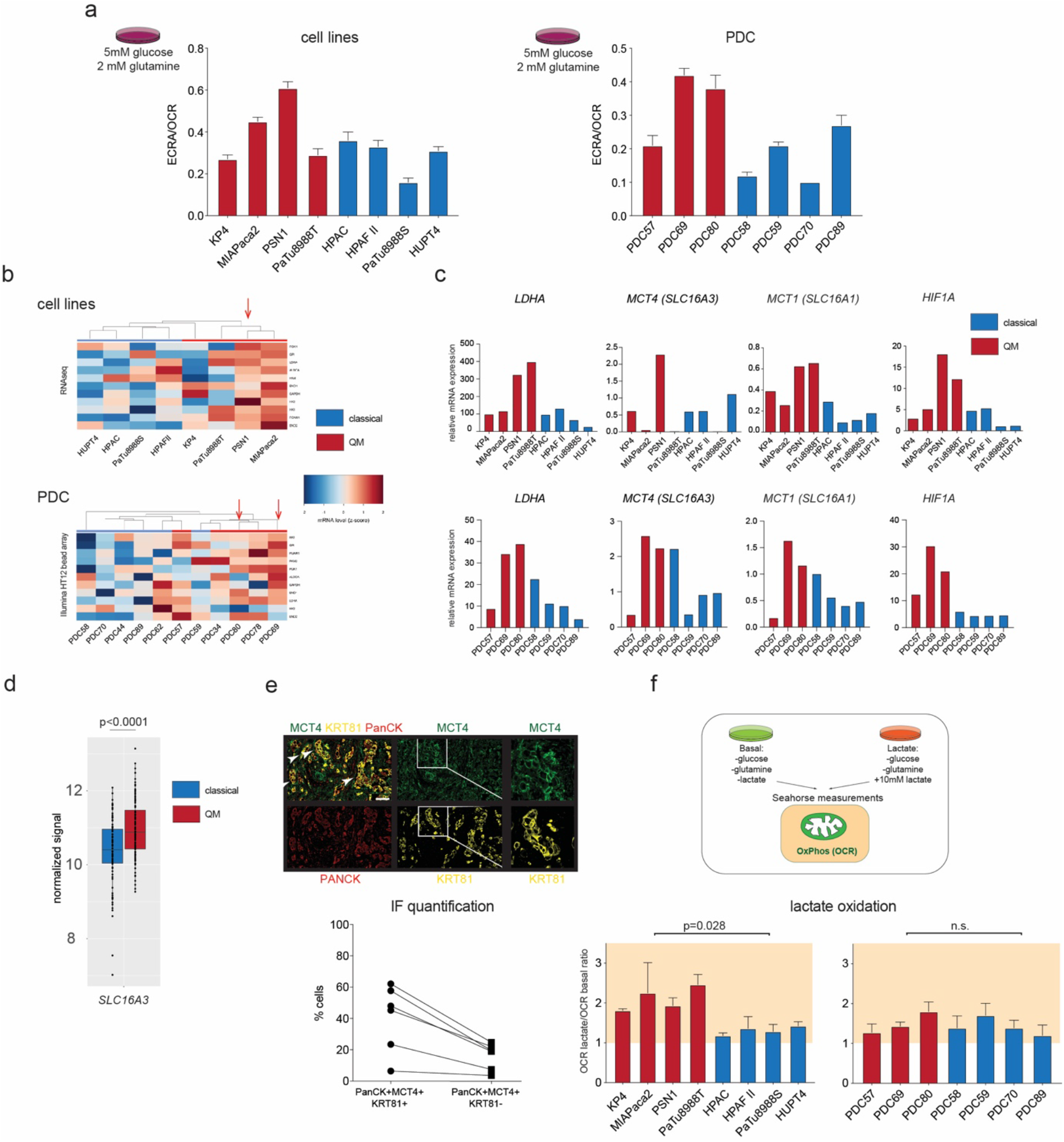
Glycolysis evaluation in PDAC. a) ECAR to OCR ratios (ECAR/OCR) as measured by seahorse metabolic flux assay for PDAC cell lines (left) and PDCs (right) cultivated in medium supplemented with 5mM glucose (physiological concentration) and 2mM glutamine. Higher ECAR/OCR ratio indicates higher glycolysis under these conditions. Presented are mean±SD values calculated for minimum of 4 wells/cell line in one experiment. b) Hierarchical clustering analysis for glycolytic genes using gene expression data for cell lines (RNA-seq) and PDCs (HT12 Illumina bead assay). Z-score: red color-high expression, blue color-low expression. PSN1, PDC69 and PDC80 show high expression of all investigated glycolytic genes. c) qPCR for *LDHA*, MCT1 (*SLC16A1*), MCT4 (*SLC16A3*) and *HIF1a* in the established and primary cells. Highest gene expression levels observed in PSN1, PDC69 and PDC80 (all QM) among cell lines and PDCs respectively. Beta-glucuronidase *(GUSB)* expression was used as house-keeper control. d) *SLC16A3* gene expression is higher in QM than in classical in patient PDAC samples. P value calculated by Student’s T-Test (unpaired, two sided). e) Multiplexed immunofluorescence staining of MCT4 (glycolysis marker), cytokeratin 81(KRT81-QM marker) and pan-cytokeratin (cancer cell marker) on 6 patient PDAC FFPE samples. White arrows indicate overlapping MCT4 and KRT81 signals. Scale bar: 10μm. Lower graph: quantification of respective populations in 6 PDAC samples by Halo. Around 30-50% of KRT81 positive cancer cells are also MCT4 positive; among KRT81 negative cancer cells, less than 20% are also positive for MCT4. Populations determined in the same sample, one line indicates one patient. f) Upper panel: schematic representation of the performed seahorse assay. Cells were cultivated in “basal” medium (no glucose, no glutamine) or in “basal” media supplemented with 10mM Sodium-L-Lactate (“basal+lactate”) for 7 hours in total and OCR levels are measured. Ratios among OCR values measured for “basal+lactate” and “basal” only media are calculated and presented. Ratio above 1 indicates increase in OCR due to lactate application. Presented are mean values of minimum 2 independent experiments (mean±SD). P values calculated by the Mann-Whitney test for QM vs classical cells.

Hierarchical clustering of transcriptome data revealed generally higher expression of glycolytic genes in QM cell lines and PDCs, especially in PSN1 and PDC69 and PDC80 (figure 3b). Notably, genes coding the glycolytic enzyme lactate dehydrogenase A (*LDHA*), lactate exporter MCT4 (*SLC16A3*) and importer MCT1(*SLC16A1*) and HIF1a, a central transcriptional and cellular regulator of hypoxia and glycolysis, were also well expressed in PSN1, PDC69 and PDC80 cells (figure 3c). MCT4 has previously been suggested to be a marker of glycolytic PDACs [Baek et al., 2014]. We also observed both in PDX and bulk PDAC tissue samples that MCT4 *(SLC16A3)* was significantly higher expressed than MCT1 *(SLC16A1)* (supplementary figure 3a), further supporting a lead role of MCT4 as lactate transporter in tissue context. Furthermore, in PDAC patient samples, MCT4 gene expression was significantly higher in QM than in classical PDACs (figure 3d)

An immunohistochemical analysis of MCT1, MCT4 and an established QM marker KRT81 [Noll et al., 2016] in FFPE samples of 30 PDACs suggested that both MCT4 and MCT1 were expressed on cancer and stromal cells with however MCT4 more expressed on cancer cells, and MCT1 in the surrounding stroma (supplementary figure 3b). Multiplex immunofluorescence for PanCytokeratin (PanCK), KRT81 and MCT4 in 6 PDAC specimens showed that the proportion of MCT4 positive cells was much higher among KRT81 positive (30-50%) than KRT81 negative cells (< 20%) (figure 3e). Furthermore, high MCT4 gene expression also correlated with poor survival, supporting the correlation of MCT4 expression and QM subtype (supplementary figure 3c).

Taken together, active glycolysis was observed in some of QM PDAC cells and correlated well with the high MCT4 expression. Our data support the use of MCT4 as a surrogate marker of QM PDACs with activated glycolysis.

### PDAC cells actively use lactate as oxidative fuel

Active re-usage of lactate by its conversion to pyruvate and subsequent oxidation in the mitochondria has been suggested in PDAC [Hui et al., 2017]. However, whether this effect is especially attributable to lactate producing high glycolytic QM PDAC cells is still not known. Intrigued by high glycolysis and consequent high expression of lactate transporters detected in some of the PDAC cells, we also addressed lactate metabolism. To investigate this, we designed a seahorse metabolic flux assay experiment, where cells were cultivated for 7 hours in i) “basal” DMEM or RPMI media without glucose or glutamine supplementation or in ii) “basal” media supplemented with lactate (basal+10mM L-lactate). Consequently, metabolic flux measurement was performed and OCR values measured in media with and without lactate were compared. Interestingly, lactate was readily used as an oxidative fuel in cell lines of both subtypes with however more pronounced OCR increase in the QM PDAC cell lines (figure 3f). Lactate treatment led to an OCR increase in all PDCs as well, without pronounced subtype dependency (figure 3f).

To substantiate this finding, we cultivated PSN1 (QM), PaTu8988T (QM) and PaTu8988S (classical) cells in physiological DMEM medium with 5mM glucose and 2mM glutamine without media change for 24-48-72-96 hours. Glucose and lactate concentrations in the media were measured at given time points. With time, glucose concentration in the media decreased and lactate increased (0-72 hours), as expected due to glucose consumption and lactate production and accumulation. Once the glucose was consumed from the medium (approx. after 72 hours in PaTu8988T/PSN1 cells), lactate concentration in the media decreased, indicating that in absence of other resources, PDAC cells start consuming self-produced lactate (supplementary figure 3d).

In conclusion, PDAC cells, regardless of subtype, not only actively produce and excrete glycolytically produced lactate but also actively re-use it potentially as an oxidative fuel. This phenomenon was more pronounced in QM than in classical PDAC cell lines.

### Metabolic inhibitors do not show subtype specific effects in primary PDAC cells

Stratification to glycolytic and oxidative PDACs is a prerequisite for patient-tailored metabolic treatment strategies. Thus, we sought to therapeutically address the observed metabolic differences and treated PDAC cells using an anti-glycolytic and two anti-oxidative metabolic drugs: the glycolytic inhibitor GNE-140 [Boudreau et al., 2016], the mitochondrial respiratory chain inhibitor phenphormin [Boudreau et al., 2016] and TriacsinC, inhibitor of FA acylation and activation for lipid synthesis, deposition and beta-oxidation [Tang et al., 2018] (figure 4a). We followed the concentration dependent inhibition of metabolic active cells via cell titer glo assay. GNE-140 treatment indeed induced a QM subtype-specific decrease in cell viability especially in the QM cell lines, being most effective in PSN1, MIAPaca2 and PaTu8988T cells. However, PDCs were in general less sensitive to GNE-140 and the observed inhibitory effects were not subtype-dependent. Phenformin treatment induced a decrease in viability equally efficient in both QM and classical PDAC cell lines, while PDCs were rather unaffected. Triacsin C was active in all cell lines with a trend towards stronger viability inhibition in QM cells, probably by targeting accumulation of fatty acid in lipid droplets observed in these cells. The compound was also active in primary cells, however without an obvious subtype-specific effect (figure 4a). Taken together, though the LDHA inhibitor GNE-140 presented stronger efficacy against QM PDAC cell lines as expected, in the PDCs we did not observe subtype specific inhibition of cell viability with neither glycolytic nor inhibitors of oxidative metabolism.

**Figure 4:**
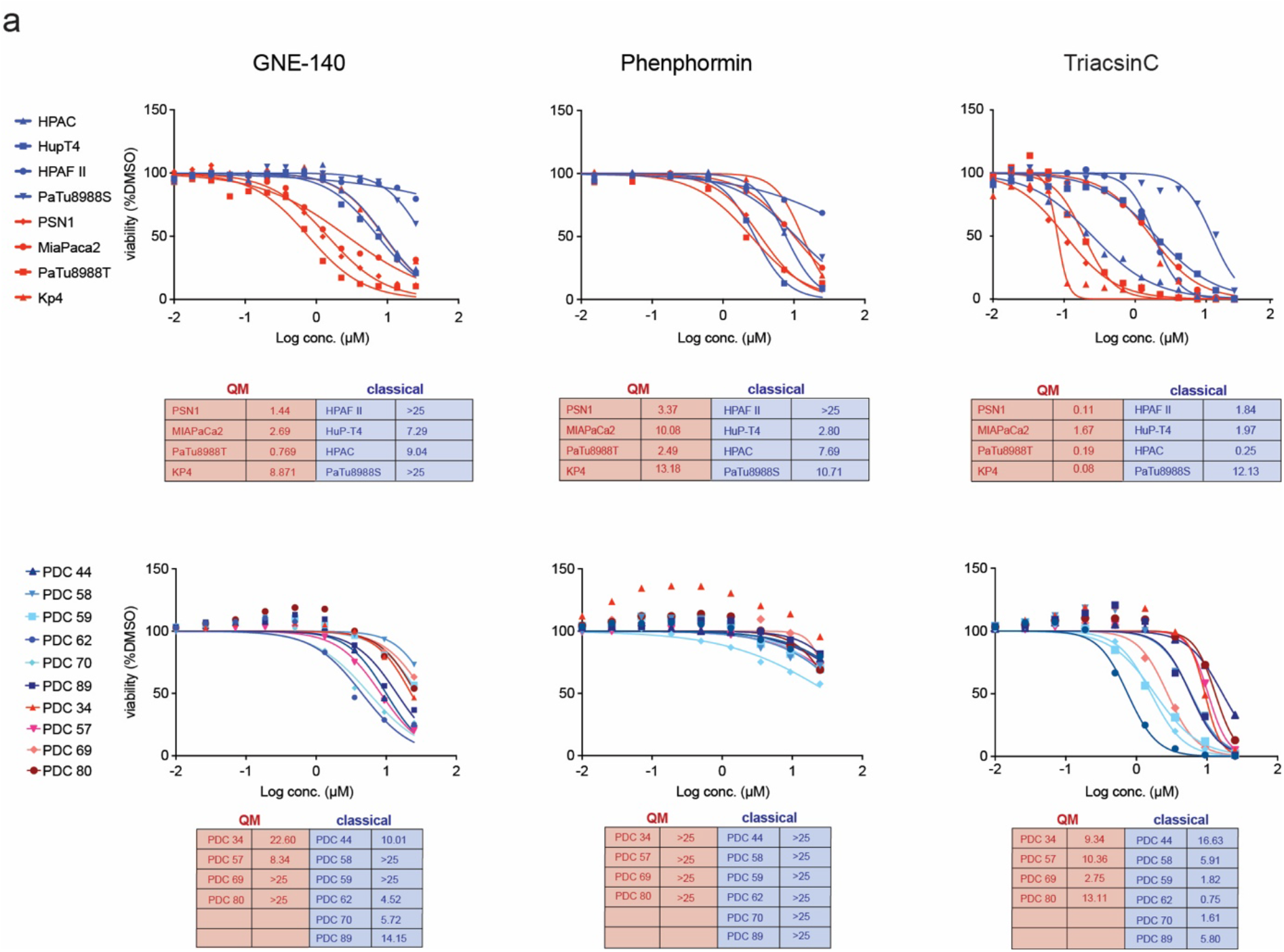
PDAC cells show differential response to metabolic inhibitors. Dose response curves of cell lines and PDCs to glycolysis inhibitor GNE-140, OxPhos inhibitor Phenphormin and lipid metabolism inhibitor TriacsinC. IC50 values are presented in tables, micromolar values (μM). Blue-classical cells, red-QM cells. GNE-140 inhibitory effects are stronger in QM than in classical PDAC cell lines. Effects in PDC lines are subtype independent. Phenphormin and TriacsinC do not show subtype specific effects in cell lines or PDCs. Presented are mean dose response curves and IC50 values of 2 independent experiments.

### Hyperpolarized magnetic resonance spectroscopy of [1-^13^C]pyruvate and [1-^13^C] lactate identifies QM tumors

Pharmacological inhibition suggested efficacy of GNE-140 in glycolytic cells arguing for the need of unequivocal identification of highly glycolytic PDACs for successful metabolic targeting. However, detection of dominant metabolic pathways driving tumor phenotypes remains a highly challenging task and is currently not established in clinical routine. Thus, we sought to explore hyperpolarized magnetic resonance spectroscopy with hyperpolarized (HP) [1-^13^C]pyruvate and [1-^13^C]lactate for potential differentiation of highly glycolytic from oxidative tumors *in vivo*. For this purpose, rats were subcutaneously implanted with glycolytic QM PSN1 and classical HPAC cells. Consistent with the respective molecular subtype, PSN1 tumors presented an undifferentiated mesenchymal phenotype, while HPAC tumors showed a more differentiated epithelial morphology (supplementary figure 4a). Once the tumors reached a minimal size of 5 x 5 mm, metabolic spectroscopy was performed. HP-[1-^13^C]pyruvate was i.v. injected into the tail vein and intratumoral distribution of HP-[1-^13^C]lactate was followed in real-time. Using MRS, significantly more HP-[1-^13^C]lactate was detected in PSN1 compared to HPAC tumors, supporting higher label exchange between pyruvate and lactate specifically in PSN1 tumors (figure 5a and 5b). Lactate dehydrogenase (LDH) enzymatic activity measured *ex vivo* after the spectroscopy experiment in snap frozen tissues was also higher in PSN1 compared to HPAC tumors (figure 5c) consistent with the *in vivo* finding.

**Figure 5:**
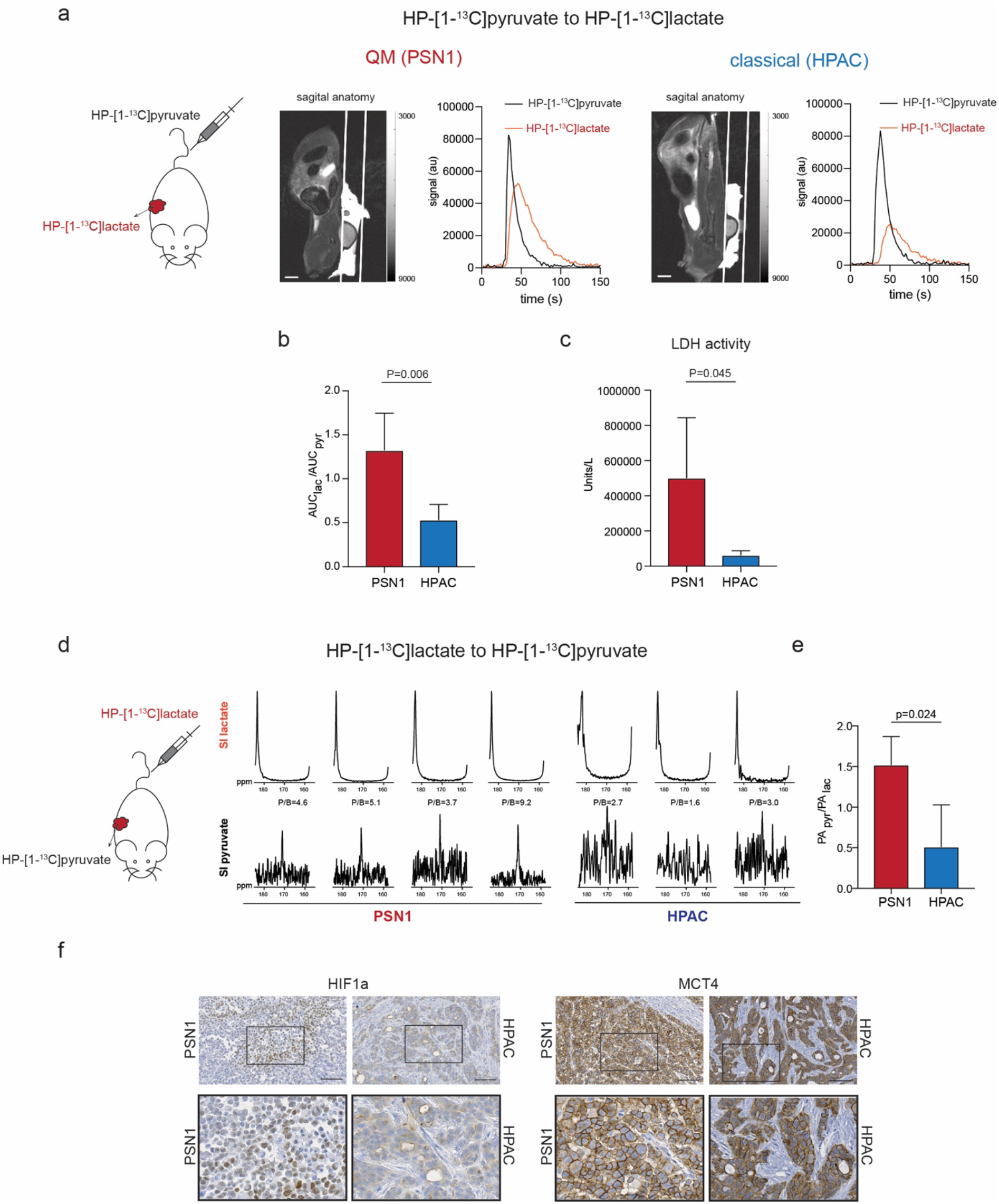
Magnetic resonance spectroscopy (MRS) of HP-[1-^13^C]pyruvate and HP-[1-^13^C]lactate inter-conversions in PSN1 (QM) and HPAC (classical) PDAC xenografts in rats. a) Left to right: schematic presentation of HP-[1-^13^C]pyruvate i.v. injection into rats with xenotransplanted PSN1 and HPAC tumors, T2-weighted sagittal anatomy image (scale bar=1cm) of a rat bearing a subcutaneous tumor and graphs demonstrating signal intensity time courses of HP-[1-^13^C]pyruvate and HP-[1-^13^C]lactate measured intratumorally in PSN1 (left) and HPAC (right) rat xenografts. The HP-[1-^13^C]lactate curve (red) is higher for PSN1 than for HPAC xenografts. b) Calculated relative AUC ratios of HP-[1-^13^C]lactate to perfused HP-[1-^13^C]pyruvate showing higher conversion rate in PSN1 (n=4; 1.325 ± 0.418) than in HPAC tumors (n=5; 0.5349 ± 0.175). P=0.006. c) *Ex vivo* measurements of lactate dehydrogenase activity in imaged tumor sample. Higher activity in PSN1 (n=5; 501794 ± 341920 U/L) than in HPAC tumors (n=5; 62796 ± 24641 U/L) detected. P=0.045. d) Left to right: schematic presentation of HP-[1-^13^C]lactate injected into rats with xenotransplanted PSN1 and HPAC tumors, signal Intensity (SI) spectra of perfused HP-[1-^13^C]lactate (top) and detected HP-[1-^13^C]pyruvate (bottom) for PSN1 (n=4) and HPAC (n=3) tumors The spectra have been summed over 10 time points covering maximum tumor enhancement and normalized to the lactate signal. Higher peak to background ratios (P/B 3.7-9.2) were observed in PSN1 tumors in comparison to P/B ratios in HPAC tumors (P/B 1.6-3.0). e) Signal intensity quantification: PApyr/PAlac ratios are significantly higher in PSN1 (1.49 ± 0.30, n=4) than in classical tumors (0.51± 0.51, n=3). P=0.024. PA-peak area. All P-values in this figure calculated by Student’s T-test (unpaired, two-sided). f) Immunohistochemistry for HIF1a and MCT4 in rat xenografts. HIF1A specific nuclear staining detected exclusively in PSN1 (QM) tumors. Scale bar=100μM.

To evaluate whether lactate can also be used by tumors *in vivo* as observed *in vitro* in seahorse experiments, we also performed the reverse experiment and injected HP-[1-^13^C]lactate in PSN1 and HPAC tumor rats *in vivo*. Intratumoral HP-[1-^13^C]pyruvate was detected in PSN1 tumors only (figure 5d) and not in HPAC tumors. Accordingly, significantly higher PApyr/PAlac ratios were measured for PSN1 than HPAC tumors (figure 5e). Taken together, highly glycolytic PSN1 xenografts could readily be discriminated based on high HP-[1-^13^C]pyruvate to HP-[1-^13^C]lactate conversion rates observed in HP-MRS. The data also showed that in glycolytic PDACs, exogenous lactate can be metabolized to pyruvate.

We further confirmed the highly glycolytic nature of PSN1 xenografts by immunohistochemical analysis of glycolytic markers HIF1A and MCT4. MCT4 showed the typical membrane-associated expression in cancer cells in both xenografts, with somewhat stronger staining intensity in PSN1 tumors (figure 5f). Intriguingly, HIF1A staining was found exclusively in the PSN1 tumors with typical nuclear expression pattern in the cancer cells (figure 5f). We also analyzed HIF1A and MCT4 expression in murine xenografts of human PDAC cell lines (supplementary figure 4b). Indeed, stronger MCT4 staining intensity was observed in the QM xenografts in general. Furthermore, specific nuclear HIF1A expression was limited to QM tumors (PSN1, KP4, MIAPaCa2, PaTu8988T), and not detected in classical tumors (HPAFII, PaTu8988S, HUPT4, HPAC) (supplementary figure 4b).

## Discussion

The challenge in PDAC is its enormous therapy resistance due to the evolution of aggressive cancer cells driven by oncogenic KRAS and loss of key tumor suppressors in a complex adapting microenvironment with various signaling effectors and biophysical and hypoxic restraints. Despite considerable genetic homogeneity with regard to oncogenic KRAS as lead driver, many studies support the existence of several molecular PDAC subtypes including classical/progenitor, QM/squamous/basal-like and hybrid states with more or less pronounced subtype specific transcriptional programs [Chan-Seng-Yue et al., 2020; Collisson et al., 2011; Moffitt et al., 2015]. Though indisputably present, functional aspects and phenotypic cues of the defined transcriptional subtypes are less well known. One key feature of PDAC is the metabolic rewiring directed by cell-autonomous and microenvironmental signals that may lead to phenotypic features not entirely captured by transcriptomic signatures. In this work, we aimed to address the functional metabolic aspects guided by transcriptome-defined classical and QM/basal like subtyping. We focused this analysis on patient-derived model systems including PDX and PDCs to value the molecular and metabolic heterogeneity in primary PDAC model systems.

Gene expression analysis in four different model systems (cell lines, PDC, PDX and bulk tissue samples) indeed identified glucose metabolism/glycolysis/hypoxia and cholesterol/lipid/fatty acid metabolism as dominating metabolic transcripts of the QM and classical subtype, respectively. This is in line with the previously observed “glycolytic” and “lipogenic” subtypes in PDAC cell lines [Daemen et al., 2015] and the recently reported “glycolytic” and “cholesterogenic” transcriptional PDAC subtypes [Karasinska et al., 2020]. In functional assays, we also observed that these transcriptional cues were correlating with metabolic behavior with however notable heterogeneity especially in patient-derived cells. Neither the lipid nor glycolytic effect was equally exposed in all of the cells of one subtype.

We identified PSN1, PDC69 and PDC80 as being typically glycolytic in seahorse assays and with high gene expression of the glycolytic markers HIF1A, LDHA and MCT4, supporting the translation of transcripts in active glucose metabolism. Interestingly, HIF1A, a major transcriptional regulator of cellular response to hypoxia [Semenza, 2010], was well expressed in highly glycolytic cells here grown in typical *in vitro* normoxic conditions, supporting intrinsic gene expression programs well preserved in QM cells. In line with our observations, MCT4 has already been suggested as marker of glycolytic PDACs with poor prognosis [Baek et al., 2014]. It should however be noted that Seahorse assays evaluate ECAR and OCR values in *in vitro* conditions and are very dependent on cell culture features such as current cellular density, growth pattern, cell cycle, current mitochondrial number [Little et al., 2020] and should be interpreted only as indication of the cellular energetic status. Better functional metabolic assays for *in vitro* and *in vivo* application are indeed needed.

Classical PDAC cells were rich in intracellular free FA, which were actively used in mitochondrial oxidation, allowing the cell to maintain the basal metabolism even in complete absence of glucose and glutamine. Our observations of different lipid/fatty acid usage of the subtypes may open a new road of further non-invasive imaging-based stratification of PDAC e.g. by using ^1^H-based diffusion-weighted magnetic resonance spectroscopy [Weidlich et al., 2019] or other quantitative MRS methods [Nemeth et al., 2018].

Heterogeneity was also present in reaction to metabolic therapies. Especially in the primary lines, neither the glycolysis inhibitor GNE-140 nor the OxPhos inhibitor Phenphormin or lipid metabolism inhibitor TriacsinC showed subtype specific effects. These results suggests that rigid classification of PDAC subtypes may not be sufficient as the basis for decisions regarding metabolic targeting approaches. Rather, individual PDACs may often present a continuum of different metabolic states that are more or less phenotypically presented depending on various cell-autonomous and non-cell-autonomous cues. Hybrid PDAC subtypes with transcriptomic signatures in between the classical and QM/basal-like states have been highlighted recently [Chan-Seng-Yue et al., 2020; Karasinska et al., 2020]. Similar to our study, a correlation of functional (seahorse) and molecular (RNA and protein) OxPhos was recently reported for PDAC cells [Masoud et al., 2020]. The authors also reported on metabolic heterogeneity and flexibility and shifts from OxPhos or glycolysis when necessary, supporting the existence of plastic metabolic states depending on the environmental challenges. It is reasonable to assume that among PDAC cells a whole spectrum from weak to highly mesenchymal and glycolytic QM, and weak to highly epithelial and lipogenic PDAC cells exists. The exclusive dependency on the one or the other metabolic pathway is thus an unlikely scenario. However, individual tumors with high activity of specific metabolic pathway may exist and their identification will be key to successful targeting. We show here that glycolytic PSN1 tumors were readily detectable with HP-MRS due to higher ^13^C-label exchange among pyruvate and lactate, indicating high activity of the last glycolytic enzyme LDHA and high intratumoral pyruvate to lactate conversion. Similarly, in breast cancer patients, high HP-[1-^13^C]pyruvate to HP-[1-^13^C]lactate conversion rates identified strongly glycolytic aggressive triple negative breast cancer with high HIF1a and MCT1 tissue expression [Gallagher et al., 2020]. This approach is already being used in personalized therapy monitoring in prostate and breast cancer [Aggarwal et al., 2017; Park et al., 2018].

We were also able to in vivo confirm the reverse effect as well, the active import and conversion of HP-[1-^13^C]lactate into HP-[1-^13^C]pyruvate in PSN1 QM-type but not in HPAC classical-type xenografts. Lactate is since recently considered as one of the important actors in tumor metabolism [Brooks, 2018]. Tumors use the advantage of lactate being the second most abundant metabolite in the systemic circulation and readily feed the TCA cycle with pyruvate generated from lactate [Faubert et al., 2017; Hui et al., 2017]. Indeed, we also observed OxPhos activation with lactate in PDAC cells, especially in the QM cell lines. It should be however noted that we performed the lactate supplementation assay in starved medium in absence of glutamine and glucose, an important TCA cycle fuel [Son et al., 2013]. Under these conditions, cells might divert to more drastic fueling of TCA cycle with lactate than physiologically typical. We speculate that the hypoxic microenvironment of the tumor favors the epithelial to mesenchymal transformation (EMT) of the cancer cells and appearance of the glycolytic QM tumors. These tumors potentially adapted their oxidative metabolism to fuels which are then locally produced, either by themselves or by neighboring cancer, stromal or immune cells.

Although HP-MRS experiments were performed on a limited number of animals, they provide evidence for the concept that PDACs with high reliance on glycolysis are potentially detectable via HP-[1-^13^C]pyruvate/lactate spectroscopy also in clinical practice. Thus, identification of highly glycolytic, aggressive PDACs by HP-[1-^13^C]pyruvate and HP-[1-^13^C]lactate spectroscopy may be used to guide and monitor tumor treatment with anti-glycolytic therapies. In contrast to biopsy-based tumor characterization, metabolic imaging allows dynamic evaluation of the whole tumor limiting sampling bias and addressing tumor heterogeneity [Hayashi et al., 2020]. Though likely not all QM tumors are potentially extremely glycolytic, non-invasive detection of highly glycolytic PDACs detected by HP-[1-^13^C]pyruvate/lactate MRS may be first candidates for successful individual metabolic targeting approaches.

## Material and methods

### Cell culture

#### PDAC cell lines

All PDAC cell lines have been obtained from the ATCC and regularly externally authenticated (at least once a year). PDAC cell lines (Psn1, Kp4, PaTu8988T, MiaPaca2, PaTu8988S, HPAC, HPAFII, HupT4) were grown in Dulbecco’s Modified Eagle Medium (DMEM, #11966025 and #A1443001, Thermo Fisher Scientific, Waltham, USA) adapted to final concentrations of 5 mM D-glucose (Thermo Fisher Scientific, Waltham, USA), 2 mM L-glutamine, 5% v/v fetal bovine serum (FBS, Thermo Fisher Scientific, Waltham, USA), and 1% v/v penicillin/streptomycin (P/S, Thermo Fisher Scientific, Waltham, USA) if not stated ootherwise.

#### Patient Derived Cells (PDCs)

For all metabolic analysis, PDC cell lines were cultivated in a 1:1 mixture of Keratinocyte-SF medium (#17005075, Thermo Fisher Scientific, Waltham, USA) and RPMI 1640 (#11879020, Thermo Fisher Scientific, Waltham, USA) adapted to final concentrations of 5mM D-glucose, 4.5mM L-glutamine, 0.26mM sodium pyruvate, and 6%v/vFBS, and 1% v/v penicillin/streptomycin (P/S, Thermo Fisher Scientific, Waltham, USA) if not stated otherwise.

#### PDX samples preparation

Establishment of the PDX mouse model was performed using surgically resected PDAC tissues collected from patients.

#### Seahorse metabolic flux assays

All assays were performed following the manufacturer’s instructions (Agilent Technologies).

#### Immunohistochemistry (IHC) and immunofluorescence

Immunohistochemistry was performed according to standard laboratory procedures on PFA fixed, FFPE tissue samples. Antibodies used in this study: MCT4, Atlas Antibodies (HPA021451); HIF1a, BD Transduction laboratories (610959); MCT1, Abcam, ab85021; KRT81, Santa Cruz, sc-100929; panCytokeratin, Abcam (ab6401);

### Hyperpolarized Magnetic Resonance Spectroscopy (HP-MRS)

#### Animal handling

All experiments were carried out in adherence to pertinent laws and regulations.

Detailed explanations of all experimental procedures can be found in supplementary material and methods section.

