## Supplementary Figures and material and methods for "Functional metabolic phenotyping of human pancreatic ductal adenocarcinoma"

Supplementary figure 1:

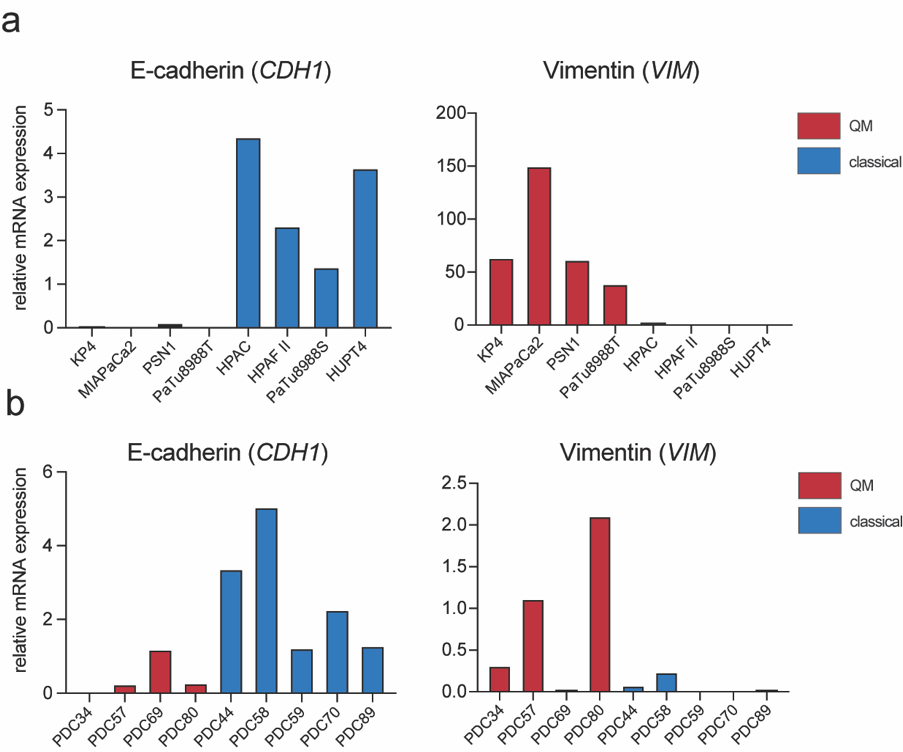

**Supplementary figure 1: Expression of E-cadherin (*CDH1*) and Vimentin (*VIM*) in PDAC cells.** qPCR analysis of epithelial marker E-cadherin (*CDH1*) and mesenchymal marker Vimentin (*VIM*) in classical and QM PDAC cells. a) PDAC cell lines b) PDCs. QM cells generally present higher expression of Vimentin (red bars) while classical cells show higher expression of E-cadherin (blue bars).

**Supplementary figure 2:**

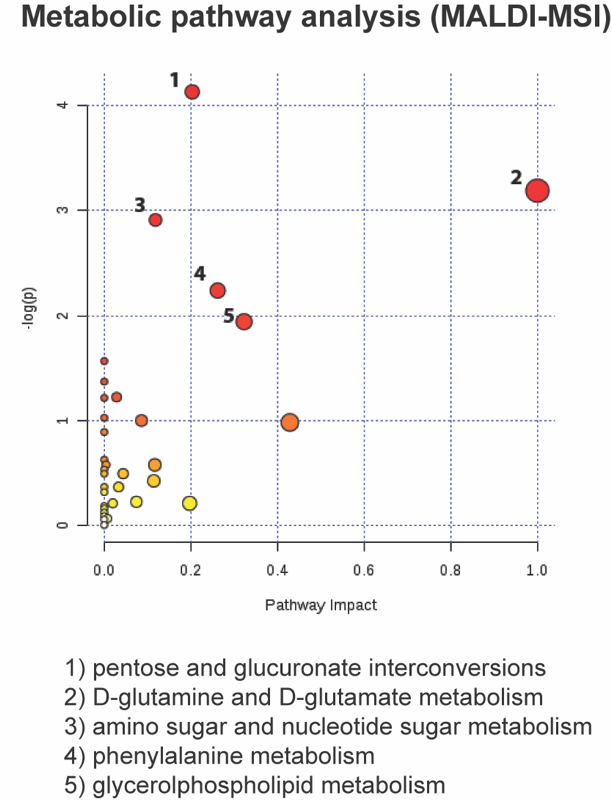

**Supplementary figure 2: Metabolic pathways analysis in PDX tissues.** Non-targeted metabolic profiling of cryo-preserved PDX tissues with MALDI-MSI (matrix-assisted laser desorption/ionization- mass spectrometry imaging) in 10 PDX PDACs (5QM vs 5 classical). Considering significantly altered metabolites between classical and QM revealed by an U-test, metabolic pathway analysis was performed. Pathways significantly changed among classical and QM samples are identified and presented on the graph (red dots).

**Supplementary figure 3:**

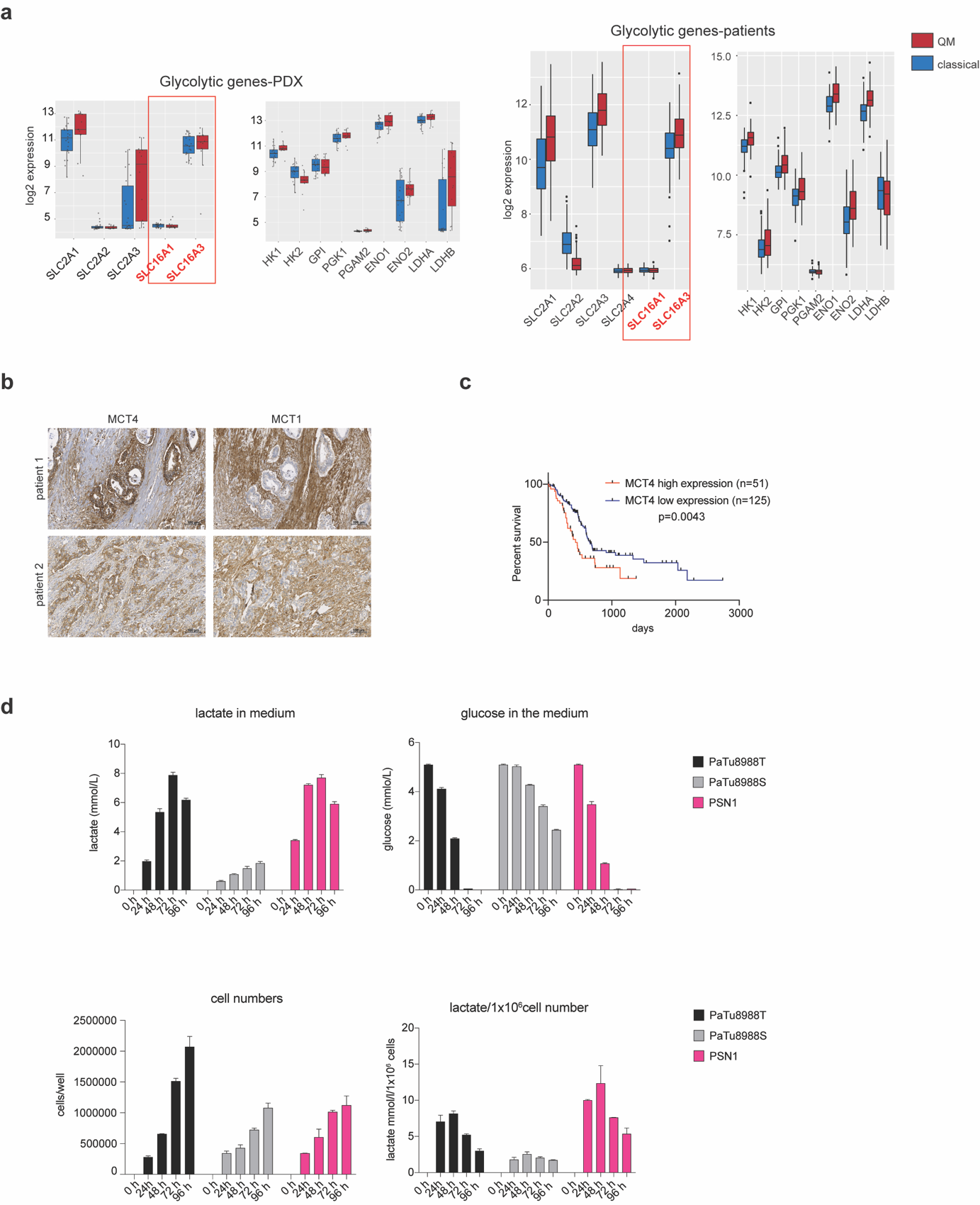
**­**

**Supplementary figure 3: Glycolysis and lactate metabolism in PDAC.** a) Gene expression analysis for glycolytic genes in PDX and patient PDAC samples (Illumina HT12 bead-array). b) Exemplarily immunohistochemistry for MCT1 and MCT4 on human PDAC FFPE samples. Both markers expressed on cancer and stromal cells with MCT4 expressed more on cancer cells and MCT1 on stromal cells. Scale bar 100µm. c) Survival analysis of PDAC patients according to *SLC16A3* (MCT4) expression. Data collected from [www.proteinatlas.org](http://www.proteinatlas.org); Patients with high MCT4 expression have worse survival. d) Measurement of lactate concentrations in the media. PSN1(QM), PaTu8988T (QM) and PaTu8988S (classical) PDAC cells were cultivated in DMEM medium (5mM glucose, 2 mM glutamine, 5% FBS) for 4 days. At defined time points an aliquot of media was collected and lactate and glucose concentration was measured. In PSN1 and PaTu8988T cells, lactate concentrations first increase (48-72h) corresponding to initial secretion, and then decrease (72h) indicating consumption of locally produced lactate once glucose is deprived. Effects less pronounced in classical PaTu8988S cells due to slower consumption of glucose. Absolute measured (mmol/l) and cell number normalized (mmol/l/1 million cells) lactate concentrations are presented.

**Supplementary figure 4:**

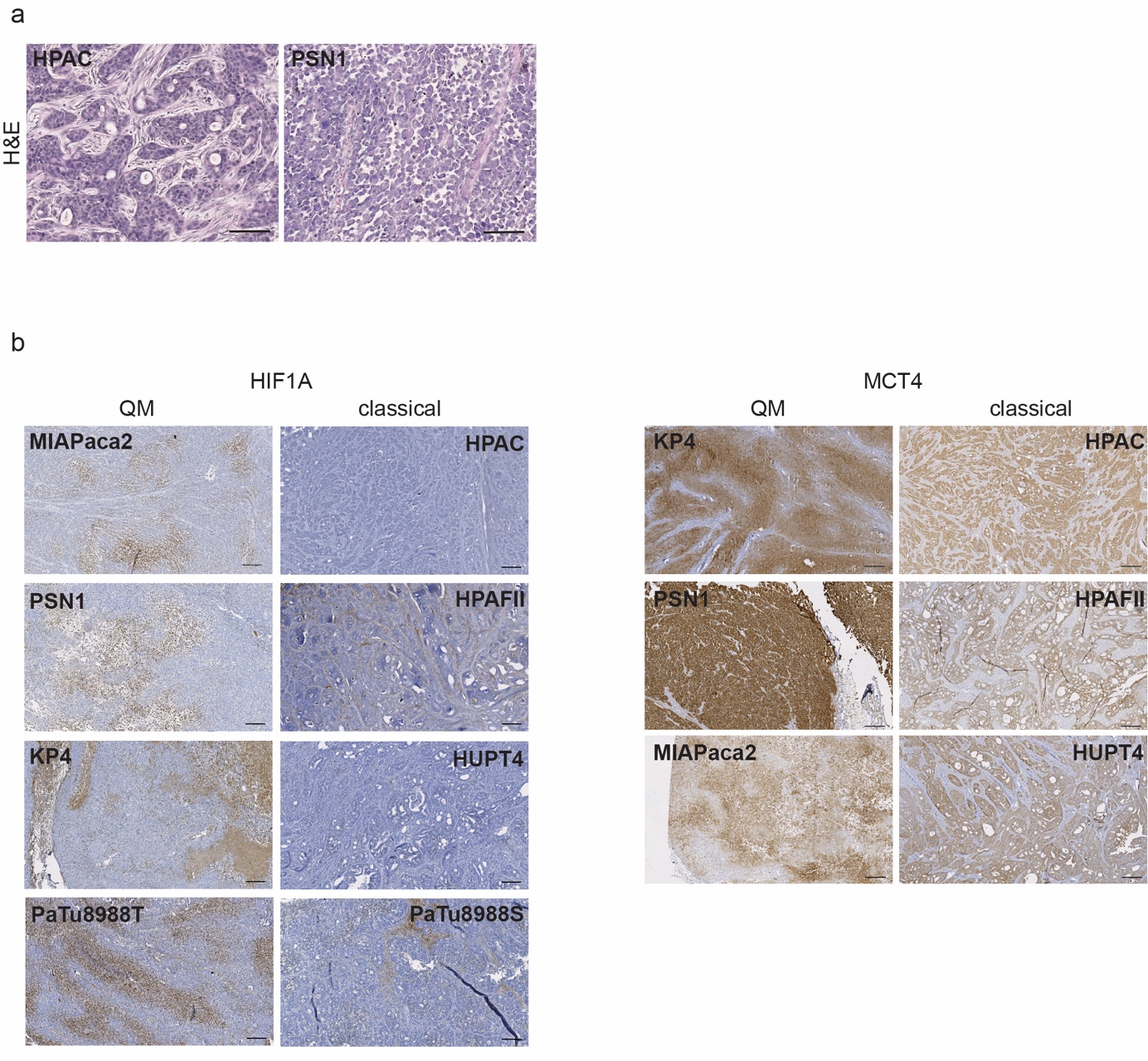

**Supplementary figure 4: Immunohistochemistry for MCT4 and HIF1a in rat and murine xenotransplanted PDAC tumors.** a) H&E histological staining of PSN1 and HPAC xenotransplanted tumors used for ^13^C-hyperpolarized HP-MRS in rats. Undifferentiated PSN1 tumors and rather glandular, differentiated HPAC tumors. b) Immunohistochemistry for HIF1A and MCT4 on murine xenotransplanted tumor tissues of human PDAC cells lines. HIF1A nuclear staining (brown horse radish peroxidase-diaminobenzidine (HRP-DAB) signals) in tumor cells is detectable only in QM xenografts (PSN1, MIAPaca2, PaTu8988T and Kp4). Some HRP-DAB positive signals are detected in the stroma of PaTu8988S and HPAF II xenografts, probably due to non-specific binding of mouse generated HIF1A antibody to murine stroma. Specific membrane-associated MCT4 positive signals are detectable in all tumors, with somewhat stronger signals intensities in PSN1, MIAPaca2 and KP4 QM tumors. Scale bar-100µM.

**Supplementary table 1:**

List of genes used as assigners for the Collisson QM and Bailey squamous subtype (Gene Program 2) used for GSEA analysis **.**

**Supplementary table 2:**

Gene Set Enrichment Analysis-list of all enriched gene sets in established PDAC cells, primary cell, PDX samples and patient cohort. Normalized Enrichment Score (NES) >1.5 and False Discovery Rate (FDR) <0.06 are used as guidelines.

**Supplementary table 3:**

List of 14 genes used for the generation of fatty acid oxidation gene set.

**Supplementary table 4:**

List of all metabolite concentrations (pmol/µg protein) in established and primary PDAC cells as measured by Biocrates absoluteID p180 kit. List of abbreviations of all metabolites also included.

**Supplementary table 5:**

Seeding densities of cells in seahorse experiments.

**Supplementary material and methods**

**Cell culture**

**PDAC cell lines**

All PDAC cell lines have been obtained from the ATCC and regularly externally authenticated by Multiplexion (at least once a year). PDAC cell lines (Psn1, Kp4, PaTu8988T, MiaPaca2, PaTu8988S, HPAC, HPAFII, HupT4) were grown in Dulbecco’s Modified Eagle Medium (DMEM, #11966025 and #A1443001, Thermo Fisher Scientific, Waltham, USA) adapted to final concentrations of 5 mM D-glucose (Thermo Fisher Scientific, Waltham, USA), 2 mM L‑glutamine, 5% v/v fetal bovine serum (FBS, Thermo Fisher Scientific, Waltham, USA), and 1% v/v penicillin/streptomycin (P/S, Thermo Fisher Scientific, Waltham, USA) if not stated otherwise. We termed this medium “low glucose DMEM”. The cells were incubated at 37°C and 5% CO_2_, provided with fresh medium every 2-3 days, and passaged at a confluency of 80‑90%.

### Patient Derived Cells (PDCs)

From 11 PDX samples, we were able to isolate and cultivate primary patient derived cells (PDCs) for further analysis. After explantation, tumor tissue was stored in RPMI (Thermo Fisher Scientific, Waltham, USA) without any supplements on ice. The tumor tissue was then minced on ice into 1-2 mm³ cubes. The minced tissue was incubated at 37°C in digestion solution (RPMI containing 5 mg/mL Collagenase II (Thermo Fisher Scientific, Waltham, USA) and 1.25 mg/mL dispase (Thermo Fisher Scientific, Waltham, USA) with agitation to dissociate the tumor tissue. Cell suspension was subsequently filtered through a 100 µm mesh and cells were collected by centrifugation (300 x g, 5 min) at room temperature. The cell pellet was washed with media and cultivated in growth media consisting of a 1:2 mixture of Keratinocyte-SF medium and RPMI 1640 (Thermo Fisher Scientific, Waltham, USA). Cells were cultured on collagen coated dishes (Corning B.V. Life Science, Amsterdam, Netherlands) in a humidified incubator at 37 °C with 5% CO2. To obtain pure PDAC cell line, cells were treated with differential trypsinization until no contaminating fibroblasts were detected by visual inspection under the microscope. The human character of the lines was confirmed by STR analysis. However, complete absence of murine cells from the PDC cultures cannot be guaranteed. Established cells were then further cultivated on standard cell culture dishes.

For all metabolic analysis, PDC cell lines were further cultivated in a 1:1 mixture of Keratinocyte-SF medium (#17005075, Thermo Fisher Scientific, Waltham, USA) and RPMI 1640 (#11879020, Thermo Fisher Scientific, Waltham, USA) adapted to final concentrations of 5mM D-glucose, 4.5mM L-glutamine, 0.26mM sodium pyruvate, and 6%v/vFBS, and 1% v/v penicillin/streptomycin (P/S, Thermo Fisher Scientific, Waltham, USA) if not stated otherwise. The cells were incubated at 37°C and 5% CO_2_, provided with fresh medium every 2-3 days, and passaged at a confluency of $\sim$90%. The primary cells were usually used 6-18 passages after thawing.

**PDX samples preparation**

Establishment of the PDX mouse model was performed using surgically resected PDAC tissues collected from patients at the Ruhr-University Bochum Comprehensive Cancer Center. Informed and written consent was obtained from all patients. The study was approved by the ethics committee of the Ruhr University Bochum (permission no. 3534-9, 3841-10, 16-5792). Patient tumor tissues were xenografted in both flanks of nude mice and expanded, isolated and re-implanted for at least three generations. All animal experiments were performed according to the guidelines of the local Animal Use and Care Committees at the Ruhr University Bochum (8.87-50.10.32.09.018, 84-02.04.2012.A328  and 81-02.04.2017.A423).

**RNA isolation and gene expression analysis**

Established/PDC cells were cultivated for 48 hours in the respective “low glucose” media. At confluence of 70-80%, cells were placed on ice and washed twice with ice-cold PBS, mechanically scratched from the plate in 1ml of ice-cold PBS and centrifuged at 4^⭘^C/400g for 5 minutes. Pelleted cells were stored in -80^⭘^C till all cells were collected for RNA isolation. RNA was isolated using the Maxwell RSC simplyRNA Cells Kit (#AS1390, Promega, Germany). Cell RNA isolation kit according to the manufacturer’s instructions. Total RNA was stored at 80^⭘^C till further processing and gene expression analysis. For PDX samples, RNA was isolated from fresh frozen PDX tumor tissue using the PARIS (Ambion) isolation kit.

For PDC and PDX samples, RNA samples were used for gene expression profiling performed using the HT12-v4 expression Bead Chip (Illumina.com) at the Genomic Core Facility at DKFZ Heidelberg. Bead intensity information was extracted using the Bioconductor package illuminio and the limma package was used for background correction. Using the necq function, quantile normalization was performed followed by log transformation. Low expression features were filtered and the probes were collapsed to the highest. The normalized expression values were then exported for downstream analysis.

For PDAC cell lines, RNA-seq analysis was performed at CeGaT GmbH. Gene expression was achieved by means of 100 bp paired end mRNA sequencing on an illumine Novaseq 600. Reads were preprocessed using casava, cutadapt and Skewer v.0.2.2 and mapped to the hg38 reference genome using STAR v.2.7.1a. Quantification was performed during alignment using STAR with the –quantMode GeneCounts parameter. A gene expression count matrix was generated with raw read counts using edgeR and stored in s DGEList object containing sample phenodata. All genes with less than 2 count per million read (cpm) in more than 20% of all samples considered were filtered out. Reads were normalized using the normalization function in edgeR and the normalized read count matrix was used for downstream analysis.

**Bioinfomatic platform for subtype determination**

For subtype determination, the normalized expression values were imported in R and the top 3000 most variable features were then selected by calculating the median absolute deviation. The selected features were centered relative to the median and unsupervised clustering was performed using the concensus clusterplus algorithm and validated using a non-negative matrix factorization approach. Gene set enrichment analysis was performed to determine the subtype of each samples by comparing with data from previously published pioneering subtyping studies [Bailey et al., 2016; Collisson et al., 2011].

To compare QM and classical mRNA data sets, Gene Set Enrichment analysis was performed using the Broad Institute GSEA software [Subramanian et al., 2005] , version 4.0.

Human PDAC gene expression data has been previously described [Jandaghi et al., 2016] (E-MTAB-1791).

Sequencing and gene expression files for cell lines, PDC and PDX samples have been uploaded to the Gene Expression Omnibus (GEO) and have following accession numbers: PDC (E-MTAB-10763); Cell lines (E-MTAB-10765); PDX (E-MTAB-10784).

**Metabolite profiling**

All cells (cell lines and PDCs) were cultivated in four 10cm round dishes in respective DMEM or RPMI media mentioned above. Media were exchanged 48h prior to collection. Cells were collected at 70-80% of confluency. Shortly, cells were seeded from the same cell suspension and in same density in four 10cm round dishes. When cells reached the desired confluency, 2 dishes were washed with room-temperature PBS (+MgCl_2_, +CaCl_2_), 1ml of ice-cold methanol was added and cells were mechanically scratched and collected in a methanol suspension that was immediately frozen and preserved in -80^⭘^C. From the remaining two dishes, cells were collected and protein concentration was measured according to the standard laboratory protocols (Pierce BCA protein assay, Thermo Fisher). Protein concentration was used for normalization of the data to assure equal loading of the assay. Metabolite profiling was performed as using the Absolute IDQ p180 kit as previously described [Charitou et al., 2019; Veyrat-Durebex et al., 2019]. Metabolite concentrations were reported in pmol/µg protein for the supplied duplicates and mean value for each cell line was calculated and used for analysis. If more than one value per metabolite was reported as being below detection range and thus extrapolated, the metabolite was excluded from analysis. If only one value was reported as extrapolated, the value and metabolite was included in the analysis. For cell lines, out of 105 measured lipid metabolites (PC-phospatidylcholines, LPC-lysophosphatidycholines, SM-spingomyelins), 84 were included in the analysis. For PDCs, 85 metabolites were included in analysis. The complete list of all included metabolites and calculated mean values is given in supplementary table 4. Volcano plots as well as heatmaps for gene expression and metabolites were generated using R software. Hierarchical clustering analysis was based on a complete linkage method with Euclidean distance. Student’s t-test (unpaired) was used for statistical analysis.

**Seahorse metabolic flux assays**

Cell lines were seeded in a seahorse 96 well microplate in 80µl final volume in respective media mentioned above and cultivated overnight in cell culture incubators (5%CO_2_, 37 C). Metabolic flux assays were performed the next day. For measurements of ECAR and OCAR levels under supplementation of 5mM glucose and 2mM glutamine, cell culture media was exchanged 1 hour prior to measurement with respective Seahorse media (Seahorse XF DMEM or Seahorse XF RPMI medium, Agilent Technologies, Santa Clara, USA) supplemented with 5mM glucose and 2mM glutamine (Agilent Technologies, Santa Clara, USA). Plates were incubated up to 1 hour in non-CO2 incubator, media was exchanged one more time shortly before the assay and then the measurement of metabolic status (ECAR and OCR) was performed using the Agilent Seahorse XFe96 machine and glycolytic rate assay protocol. Measurement 3 (basal measurements) was used for calculation of basal ECAR to OCR ratios. Attention was payed that OCR and ECAR values were are in the optimal measurement range of the instrument (ECAR 10-100, OCR 20-160) and that cells are viable (visual inspection) and around 80%-90% confluent. In all assays, 4-8 wells were seeded with cells of one cell line and used for calculation of mean value per cell line. Assays were analyzed using the Wave 2.6.0 software. In figure 3a presented are mean±SD values calculated from 4-8 wells/cell line in one experiment. Assays were repeated at least twice for cell lines and for PDCs with similar results observed.

**Basal energy measurements under glucose/glutamine starvation and lactate supplementation assay**

Cells were seeded in a Seahorse 96 well microplate coated with Cell-Tak cell adhesive in their usual respective cultivation media mentioned above (final volume: 80 µL/well). After over-night incubation in cell culture incubators (5%CO_2_, 37 C), the medium was exchanged for non-supplemented DMEM or RPMI –“basal” media (no glucose, no glutamine) or supplemented with10 mM L-lactate (sodium L-lactate, Sigma-Aldrich, 867561). Media were lacking glucose, glutamine or sodium pyruvate but were supplemented with 1% v/v FBS. The cells were incubated in the respective starved/basal (2-3 wells) or lactate supplemented (2-3 wells) medium at 37°C and 5% CO_2_ for 6 hours. Prior to the seahorse measurement, medium was changed to Seahorse XF DMEM or Seahorse XF RPMI medium (Agilent Technologies, Santa Clara, USA) either without any supplementation -“basal Seahorse” media or supplemented with or 10 mM L-lactate (sodium L-lactate, Sigma-Aldrich, 867561) -“basal Seahorse+lactate” and the cells were incubated in a non-CO_2_ incubator at 37°C for 45-60 min. For all Seahorse media the pH was adjusted to 7.4 according to the manufacturer’s instructions. 6 basal measurements of OCR and ECAR were performed, measurement 3 or 4 used for calculations. For estimation of OCR values under starvation (no glucose, no glutamine), OCR values were normalized to the 10.000 of seeded cells. In figure 2e, mean values±SD calculated from 2-3wells/cell line in one experiment are presented. The assay was repeated at least twice with similar result observed. For lactate supplementation assay, mean OCR of lactate-supplemented wells was divided with mean OCR of non-supplemented/basal wells and OCR-lactate/OCR-basal ratio was obtained for each cell line. The assays were repeated at least twice. OCR-lactate/OCR-basal ratios are calculated per cell line/per assay and figure 3f presents mean value of ratios from all performed assays.

For all seahorse assays, seeding cell densities per cell line are presented in supplementary Table 5.

**Lactate/Glucose concentrations in the media**

Changes in the lactate and glucose concentrations in culture medium of PDAC cell lines were observed over a 24-48-72-96 hour time period. The assay was performed in a 6-well dish format with three technical replicates per cell line and time point. Cells were seeded in respective DMEM media mentioned above (final volume: 1.5 mL/well). At defined time points (24-48-72-96 hours) , medium was collected (600µL) and the cells in the respective wells were tripsinized and collected and cell number determined using the TC20 automated cell counter (BIO-RAD Laboratories, Hercules, USA). Lactate and glucose concentrations in the media were determined in the central laboratory facilities at the university clinic in Essen. Measured lactate concentrations were corrected for lactate concentration present in the media without cells (0.5mmol/l) and for evaporation factor (media evaporation observed after 48-72-96 hours of cultivation). In addition, lactate concentrations were normalized to cell numbers and normalized values are presented in the figure as well.

**Quantitative Real-Time PCR**

RNA isolation was done with Maxwell® RSC simplyRNA Cells Kit (Promega, AS1390) by using Maxwell® RSC Instrument (Promega, AS4500) according to manufactrer’s instructions. Invitrogen™ SuperScript™ IV First-Strand Synthesis System (ThermoFischer Scientific, 18091050) was used to synthesize cDNA. Approximatelly 1-2µg of total RNA was used for every cDNA synthesis. Semi-quantitative PCR was performed with SYBR GREEN PCR Master Mix (Roche, 04707516001). Relative expression of genes of interests from each sample were determined with LightCycler 4800 instrument (Roche, #05015278001). Beta-glucuronidase (*GUSB*) gene were used as reference. Results were calculated using the standard curve and the comparative delta Ct method. Mann-Whitney test was used for statistical analysis. Primers sequences were as follows: GUSBfwd:TGCAGGTGATGGAAGAAGTG and GUSBrvs: TTGCTCACAAAGGTCACAGG; HIF1Afwd:GCCGCTGGAGACACAATCAT and HIF1Arvs: TGGGTGAGGGGAGCATTACA; SLC16A1fwd: CACCAGCGAAGTGTCATGGA and SLC16A1rvs: ATCAAGCCACAGCCTGACAA; SLC16A3fwd: ATCACTGGCTTCTCCTACGC and SLC16A3rvs: CTGTAGCCGATCCCAAACTC;

**Giantin (Golgi-complex) Immunofluorescence**

For Giantin staining, PDAC cells were grown in high glucose DMEM (Invitrogen, 31966021) supplemented with 5% FBS and 1% Pen/Strep. Primary cells were grown in the respective RPMI media mentioned above. For imaging experiments cells were seeded in Eppendorf/Falcon 8-well chamber slides. 48h after last media exchange, cells were fixed in ice-cold methanol for 10 min and rinsed with PBS. After blocking and permeabilization in 10% FBS in PBS-T (PBS with 0.1% Triton X-100), cells were incubated in primary antibody solution (anti-Giantin antibody (Abcam, Cat# ab37266, RRID:AB_880195) 1:200 in 1% BSA in PBS-T) o/n at 4°C. Upon incubation with secondary antibody solution (goat anti-mouse secondary antibody conjugated to Alexa Fluor 546 (Invitrogen, A-11003)) for 60 min at RT, nuclei were stained with 5 µg/ml Hoechst (Invitrogen, H1399) in PBS for 20 min. Fluorescence images were taken at a Leica TCS SP8 confocal laser scanning microscope using a 20x/NA 0.75 dry objective. Images were processed with Fiji [Schindelin et al., 2012].

**Free Fatty Acid (FFA) quantification**

Intracellular Free Fatty Acid were quantified according to manufacturer’s instructions (Abcam, ab65341). Briefly, cells were grown in their respective media mentioned above in 10cm round dishes. At 70-80% confluency, medium was changed to FBS-free medium and cells were incubated at 37˚C/ 5 % CO2 for 24 hours. Consequently, the cells were washed twice with ice-cold PBS to remove traces of media and collected by mechanical scratching in 1ml PBS. Cell count was determined TC20 automated cell counter and 1x10^6^ cells were used for the assay. Mann-Whitney test was used for statistical analysis.

**OilRedO staining for lipid droplets**

All PDAC cells were grown in their respective media mentioned above in 8-well cell culture chambers (Falcon).The staining was performed according to manufacturer’s instructions (Lipid (OilRedO) staining kit, Sigma Aldrich MAK194-1KT). Presented are inserts of images taken with 40x objective with Zeiss Axio Imager A2 Light Microscope.

**MALDI mass spectrometry imaging**

PDX tissue samples were isolated from the animals, snap frozen in liquid N_2_ and kept in -80 untill processing. Tissue samples were prepared as previously described [Aichler et al., 2017; Ly et al., 2016]. MALDI-MSI was performed using a Bruker Solarix 7T FTICR-MS (Bruker Daltonik, Bremen, Germany) in negative ion mode with a lateral resolution of 60 µm.

The acquired mass spectrometry imaging data underwent spectra processing with FlexImaging v. 4.0 software (Bruker Daltonics). The raw data were normalized against the root mean square of all data points. The average spectra of defined regions of interest were exported as .csv files and subsequently processed by MATLAB R2013a (Mathworks, Natick, MA, USA). A self- implemented MATLAB analysis pipeline was performed as previously described [Aichler et al., 2017] [Ly et al., 2016]. To identify statistically significant differences in m/z values, the peak lists were analyzed using the Wilcoxon-Mann-Whitney -test. As a result, a list of significantly different metabolites could be achieved with a corresponding p-value of ≤0.05. Metabolites were estimated by matching accurate mass with databases, as described elsewhere [Aichler et al., 2017; Ly et al., 2016]: METLIN (http://metlin.scripps.edu) [Smith et al., 2005] and the Human Metabolome Database (www.hmdb.ca) [Wishart et al., 2018]. These databases were used in conjunction with MetaboAnalyst (www.metaboanalyst.ca) [Chong et al., 2018] and KEGG (www.genome.jp/kegg)[Kanehisa and Goto, 2000] for pathway analysis, as previously shown [Sun et al., 2018].

**Cell viability assays**

Cell viability was determined using the CellTiter-Glo® (CTG) Luminescent Cell Viability Assay (Promega) according to manufacturer’s instruction. Cell numbers (number of cells seeded per well) were optimized for 96-well cell plate format using CTG assay, attention was payed that optimal cell numbers are in linear range of luminescence measurement. Metabolic inhibitors GNE-140 (MedChemExpres), TriacsinC (Cayman Chemical) and Phenphormin (Sigma-Aldrich or Cayman Chemical) were dissolved in dimethyl sulfoxide (DMSO) (Sigma-Aldrich) and printed in the indicated logarithmic concentration ranges using the D300e Digital Dispenser (Tecan). The DMSO concentration in each well was adjusted to the highest value on the plate which was set to < 0.1% of the assay volume. Sealed plates were frozen at ‑80°C until use. Cells were grown in their respective media and detached by 0.05% trypsin - ethylenediamine tetraacetic acid (EDTA) (1x) (Thermo Fisher Scientific) and recovered by centrifugation. Optimized cell numbers were seeded in 100µl of respective media with the Multidrop Combi Dispenser (Thermo Fisher Scientific) onto the pre-printed plates and incubated at 37°C and 5% CO_2_ for 72 hours. Cell viability was determined using the CellTiter-Glo® Luminescent Cell Viability Assay (Promega) according to manufacturer’s instruction. The luminescence signal was measured with a Tecan Spark® 10 M multiplate reader (Tecan). Data were normalized to the signal of DMSO treated cells. IC50 determination was performed using the Graph Pad Prism v. 7 ‘log (inhibitor) vs. normalized response (variable slope)’ equation.

**Immunohistochemistry (IHC) and immunofluorescence**

Immunohistochemistry was performed according to standard laboratory procedures on PFA fixed, FFPE tissue samples. Antibodies used in this study: MCT4, Atlas Antibodies (Sigma Aldrich, Cat#HPA021451, RRID:AB_1853663); HIF1a, BD Transduction laboratories #610959; MCT1, Abcam, #ab85021; KRT81, Santa Cruz, #sc-100929; panCytokeratin , Abcam #ab6401;. Signals were developed using horseradish peroxidase-DAB detection system (brown signals). Multiplexed IF was performed using the Opal multiplex system (NEL811001KT, Perkin Elmer, MA) according to manufacturer’s instruction. In brief, FFPE sections were deparaffinized and then fixed with 4% paraformaldehyde prior to antigen retrieval by heat-induced epitope retrieval using citrate buffer (pH 6) or Tris/EDTA (pH 9). Each section was put through several sequential rounds of staining; each includes endogenous peroxidase blocking and non-specific protein blocking, followed by primary antibody and corresponding secondary horseradish peroxidase-conjugated polymer (Zytomed Systems, Germany or Perkin Elmer). Each horseradish peroxidase-conjugated polymer mediated the covalent binding of different fluorophore using tyramide signal amplification. Such covalent reaction was followed by additional antigen retrieval in heated citric buffer (pH6) or Tris/EDTA (pH9) for 10 min to remove antibodies before the next round of staining. After all sequential staining reactions, sections were counterstained with DAPI (Vector lab). Slides were scanned and digitalized by Zeiss Axio Scanner Z.1 (Carl Zeiss AG, Germany) with 10x objective magnification. The whole-slide images were analyzed using digital image analysis software (HALO^TM^ Version 2.0, Indica Labs, Corrales, NM). Total cell number, and the number of MCT4^+^, Krt81^+^, PanCK^+^ cells were quantified in the total fraction of tissue surface area. The following subsets were defined and quantified: Krt81^+^PanCK^+^, Krt81-PanCK^+^, MCT4^+^Krt81^+^PanCK^+^, MCT4^-^Krt81^+^PanCK^+^, MCT4^+^Krt81-PanCK^+^, MCT4-Krt81-PanCK^+^.  Acellular and necrotic areas were excluded from analysis.

**Patient survival analysis**

Gene expression data for MCT4 (SLC16A3), HIF1A, CMYC and LDHA were extracted from publicly available resource [www.proteinatlas.org](http://www.proteinatlas.org) where RNA-seq data is reported as mean FPKM (TCGA) [Uhlen et al., 2017]. The best expression cutoff was accepted from the [www.proteinatlas.org](http://www.proteinatlas.org) as well as the Log-rank P-values presented on the figures.

**Hyperpolarized Magnetic Resonance Spectroscopy (HP-MRS)**

**Animal handling**

Approval of the animal protection and welfare review board was received prior to study initiation (ROB-55.2-2532.Vet_02-18-91). All experiments were carried out in adherence to pertinent laws and regulations. For all interventions rats ware anesthetized with inhalation of isoflurane 2.5% (v/v) in an oxygen flow rate of 2 l/min, breathing rates and temperature were constantly monitored and kept in standard range (50-70 breathing rate, 37-39 °C) and a tail-vein-catheter was inserted.

**Spectroscopy tumor model and tumor sample preparation**

PSN1/HPAC cells were cultured under standard condition in high glucose DMEM supplemented with 1% MEM Non-Essential Amino Acids Solution , 10% v/v fetal bovine serum and 1% v/v penicillin/streptomycin (all Thermo Fisher Scientific, Waltham, USA). and 1*10^7^ cells were implanted subcutaneously (s.c.) into the back of male or female 6 weeks-old Crl:NIH-*Foxn1^rnu^* rats (Charles River). Tumors of minimum size of 5x5x5 mm^3^ were used for HP-MRS experiments. After euthanization, tumors were removed rapidly and divided into two parts: one was snap frozen in liquid nitrogen, and the other part used for histological tumor sample evaluation.

**LDH activity assay**

Ex vivo enzyme measurements of total LDH were performed photometrically on supernatant of 100 µg frozen tumor tissue, shredded in 1 ml of RIPA buffer (1M Tris-HCl (ph 8), 1M NaCl, Nonidet P40, sodium deoxycholate, distilled water) and centrifuged (4°C; 5000 rpm; 15 min) using cobas® c 701/702 system (Roche/Hitachi) following manufacturer’s instructions in the clinical routine laboratory in the department of clinical chemistry of Technical University of Munich.

**Substrates and polarization procedure**

The hyperpolarization of pyruvate was performed as previously described [Hundshammer et al., 2018]. Briefly: 14 M [1-^13^C]pyruvate (Merck, Darmstadt, Germany) supplemented with 15 mM OX063 trityl radical (Oxford Instruments, Abingdon, UK) and 1 mM Dotarem^®^ were polarized with a HyperSense (Oxford Instruments, Abingdon, United Kingdom) for ~60 min at 1.3 K using a microwave frequency of 94.19 GHz and 100 mW power. The sample was dissolved in 3.4 ± 0.3 mL of buffered solution pressurized to 10 bar and heated to 180 °C containing 80 mM TRIS, 0.1 g/l EDTA and 80 mM sodium hydroxide (NaOH) resulting in a 80 mM [1-^13^C]pyruvate solution with mean pH of 6.9 ± 0.4. Apparent *T*_1_ values of 49.2 ± 1.5 s for hyperpolarized [1-^13^C]pyruvate in this injection solution were measured in a 43 MHz Spinsolve carbon benchtop spectrometer (Magritek, Aachen, Germany; Wellington, New Zealand) using a single pulse acquisition with flip angle 10° and TR 5 s.

The hyperpolarization of [1-^13^C]lactate was performed as described before [Park et al., 2015]. Briefly: 2.21 M of 45-55% w/w [1-^13^C]lactate (Merck, Germany) supplemented with 30% DMSO and 15 mM OX063 trityl radical were polarized using above described instruments and conditioned with a microwave frequency of 94.18 GHz and 100 mW power for ~200 min. Dissolution was performed with 3.1 ml solution of 80 mM Tris/D_2_O/1M NaOD resulting in a 100 mM [1-^13^C]lactate solution with mean pH of 7.3 ± 0.2. Apparent *T*_1_ values of 49.2 ± 2 s for hyperpolarized [1-^13^C]lactate in this injection solution were measured in a 43 MHz Spinsolve carbon benchtop spectrometer (Magritek, Aachen, Germany; Wellington, New Zealand) using a single pulse acquisition with flip angle 10° and TR 5 s.

**MRI Imaging**

All MRI experiments were performed with a small animal 7 T preclinical scanner (Agilent/GE magnet, Bruker AVANCE III HD electronics) with a dual-tuned ^1^H^1^/^13^C volume resonator with inner diameter 72 mm for anatomical proton imaging and ^13^C excitation, and 20 mm diameter ^13^C flexible surface receive coils (RAPID Biomedical). Surface receive coils were placed of top of the s.c. tumors that were covered in Carbopol® 980 (Caesar & Loretz GmbH, Germany) to reduce B_0_ field shimming artefacts. Initial proton localizer images were used to adjust the animal position so that tumors were placed at the center of the magnet and volume resonator. Anatomical proton MR images were acquired using a multi-slice T2-weighted RARE (rapid acquisition with relaxation enhancement) sequence. Axial, coronal, and sagittal images were acquired in order to guide placement of the ^13^C spectroscopy slice on the tumor, without involvement of adjacent tissue. The imaging parameters of the sagittal T2-weighted scans were echo time TE = 30 ms, repetition time TR = 3 s, field of view 128 x 72 mm^2^, in-plane resolution 0.25 x 0.25 mm^2^, 25 slices of 2 mm thickness, and 3 image averages.

Pyruvate-lactate metabolism was measured with multi-frame slice spectroscopy (MRS, 15 mm slice thickness) using alternating metabolite-frequency-selective excitation (flip angle 30°, 250 Hz transmit bandwidth, both metabolites separately excited and measured every 2 s) while injecting hyperpolarized HP-[1-^13^C]pyruvate or HP-[1-^13^C]lactate. Procedure optimization is described in detail in our previous study, which used two of the same animals bearing PSN1 tumors that were used in this study as well [Topping et al., 2021].

For data analysis after HP-[1-^13^C]Pyruvate injection, lactate and pyruvate spectral peak heights were summed over all time points after signal appeared. The ratio of these areas under curves (AUC_lac_/AUC_pyr_) gives a marker sensitive to conversion of pyruvate to lactate [Hill et al., 2013].

For data analysis after HP-[1-^13^C]Lactate injection, due to low pyruvate signal, the signal intensities of lactate and pyruvate spectra were averaged over 10 time points near the maximum intensity and then fit with a constant offset plus a Lorentzian function with fixed 30 Hz full-width at half-maximum to determine the peak area (PA), using the least-squares curve fit function in MatLab. A ratio of PA_pyr_/PA_lac_ was used to describe metabolic differences between the groups. Peak to background (P/B) ratios were calculated as peak value divided by mean signal value of the first 200 points. All statistical analyses for the imaging were performed using Prism 7 (GraphPad Software).

**Statistical analysis**

All used tests and p-values are indicated on figures.
